# Phylogenetic barriers to horizontal transfer of antimicrobial peptide resistance genes in the human gut microbiota

**DOI:** 10.1101/385831

**Authors:** Bálint Kintses, Orsolya Méhi, Eszter Ari, Mónika Számel, Ádám Györkei, Pramod K. Jangir, István Nagy, Ferenc Pál, Gergö Fekete, Roland Tengölics, Ákos Nyerges, István Likó, Balázs Bálint, Bálint Márk Vásárhelyi, Misshelle Bustamante, Balázs Papp, Csaba Pál

## Abstract

The human gut microbiota has adapted to the presence of antimicrobial peptides (AMPs) that are ancient components of immune defence. Despite important medical relevance, it has remained unclear whether AMP resistance genes in the gut microbiome are available for genetic exchange between bacterial species. Here we show that AMP- and antibiotic-resistance genes differ in their mobilization patterns and functional compatibilities with new bacterial hosts. First, whereas AMP resistance genes are widespread in the gut microbiome, their rate of horizontal transfer is lower than that of antibiotic resistance genes. Second, gut microbiota culturing and functional metagenomics revealed that AMP resistance genes originating from phylogenetically distant bacteria only have a limited potential to confer resistance in *Escherichia coli*, an intrinsically susceptible species. Third, the phenotypic impact of acquired AMP resistance genes heavily depends on the genetic background of the recipient bacteria. Taken together, functional compatibility with the new bacterial host emerges as a key factor limiting the genetic exchange of AMP resistance genes. Finally, our results suggest that AMPs induce highly specific changes in the composition of the human microbiota with implications for disease risks.

## Introduction

The maintenance of homeostasis between the gut microbiota and the human host tissues entails a complex co-evolutionary relationship^1,2^. Specialized cells in the intestinal epithelium restrict microbes to the lumen, control the composition of commensal inhabitants, and ensure removal of pathogens^3,4^. Cationic host antimicrobial peptides (AMPs) have crucial roles in this process^5^. They are among the most ancient and efficient components of the innate immune defence in multicellular organisms and have retained their efficacy for millions of years^5,6^. As AMPs have a broad spectrum of activity, much effort has been put into finding potential novel antibacterial drugs among AMPs ^7,8^.

However, therapeutic use of AMPs may drive bacterial evolution of resistance to our own immunity peptides ^9,10^. Therefore, it is of central importance to establish whether AMP resistance genes in the gut microbiome are available for genetic exchange with other bacterial species.

Several lines of observation support the plausibility of this scenario. The gut bacterial community is a rich source of mobile antibiotic resistance genes^11^, and certain abundant gut bacterial species exhibit high levels of intrinsic resistance to AMPs^12^. Moreover, even single genes can confer high AMP resistance in Bacteroidetes^12^. However, beyond the recent discovery of a horizontally spreading resistance gene family^13,14^, the mobility of AMP resistance-encoding genes across bacterial species has remained a *terra incognita*.

Here, we applied an integrated approach to systematically characterize the mobilization potential of the AMP resistance gene reservoir in the human gut microbiome. First, we examined the patterns of horizontal gene transfer events involving AMP resistance genes by analyzing bacterial genome sequences from the human gut. Next, we experimentally probed the functional compatibility of these AMP resistance genes with a susceptible host, *E. coli*, by performing functional metagenomic selections in the presence of diverse AMPs. By comparing these results with those obtained for a set of clinically relevant small-molecule antibiotics with well-characterized resistomes, we found that AMP resistance genes are less frequently mobilized and have a lower potential to confer resistance in a phylogenetically distant new host. Finally, we demonstrated that the phenotypic impact of acquiring AMP resistance genes frequently depends on the host’s genetic background. Overall, these findings indicate that lack of functional compatibility of AMP resistance genes with new bacterial hosts limits their mobility in the gut microbiota.

## Results

### Infrequent horizontal transfer of AMP resistance genes in the gut microbiota

We begin by asking whether the genetic determinants of resistance to AMPs and antibiotics, respectively, differ in their rate of horizontal transfer in the human gut microbiota. To systematically address this issue, we first collected a comprehensive set of previously characterized AMP- and antibiotic-resistance genes from literature and databases, yielding a comprehensive catalogue of 105 and 200 AMP- and antibiotic-resistance gene families, respectively (see Methods and Table S1). Next, we compared the frequencies of these previously identified resistance genes in a catalogue of 37,853 horizontally transferred genes from 567 genome sequences of phylogenetically diverse bacterial species in the human gut microbiota^15^. This mobile gene catalogue relies on the identification of nearly identical genes that are shared by distantly related bacterial genomes and thereby provides a snapshot on the gene set subjected to recent horizontal gene transfer events in a representative sample of the human gut microbiome^15^. We identified homologs of the literature-curated resistance genes for which at least one transfer event was reported (i.e., those present in the mobile gene pool; see Methods and Table S2).

We found that the relative frequency of AMP resistance genes within the pool of mobile genes was 6-fold lower than that of antibiotic resistance genes, in spite of their similar frequencies in the genomes of the gut microbiota (Figure 1A, Table S2). Moreover, the unique transferred genes were shared between fewer bacterial species, indicating fewer transfer events per gene (Figure 1B). Overall, these results suggest that AMP resistance genes are less frequently transferred across bacterial species in the human gut.

**Figure 1.**
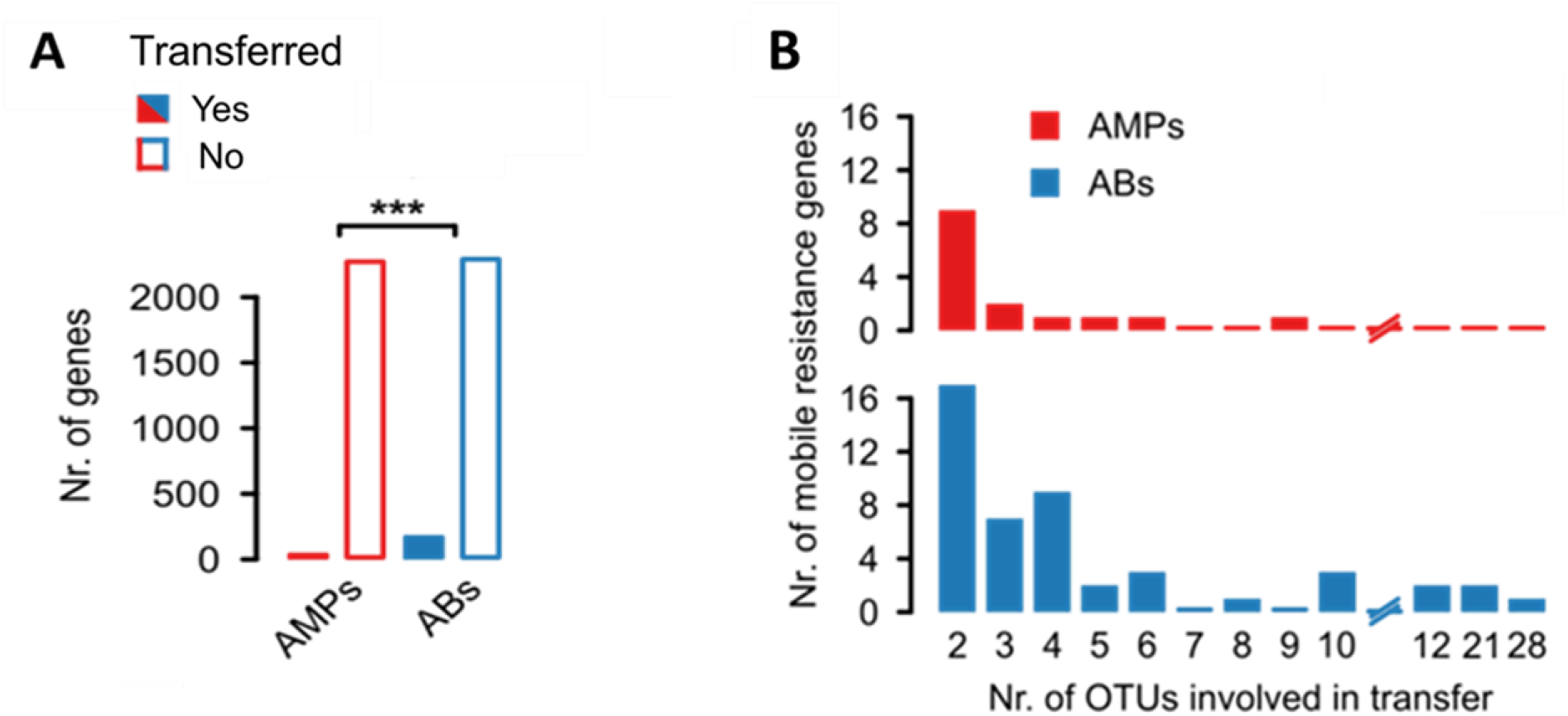
AMP resistance genes are less frequently transferred in the human gut microbiome than antibiotic resistance genes. **A)** The number of the known AMP-(red bars) and antibiotic-resistance genes (blue bars) from the gut microbiome that are transferred (filled bars) / not transferred (empty bars). Known resistance genes were identified using blast sequence similarity searches (see Methods). *** indicates significant difference (P = 10^−24^, two-tailed Fisher’s exact test, *n*=4714). **B)** Unique mobile AMP resistance genes (red bars) were involved in only approximately half as many between-species transfer events as antibiotic resistance genes (P=0.03, two-sided negative binomial regression, n=45 (AMPs) and *n*=251 (ABs)).

### Short genomic fragments from the gut microbiota rarely confer AMP resistance

One possible reason for the low mobilization of AMP resistance genes could be that AMP resistance is an intrinsic property of certain bacteria shaped by multi-gene networks^16^. Genes involved in AMP resistance may display strong epistatic interactions, and therefore they may have little or no impact on resistance individually. If it was so, horizontal gene transfer of single genes or transcriptional units encoded by short genomic fragments would not provide resistance in the recipient bacterial species. Indeed, AMPs interact with the cell membrane, a highly interconnected cellular structure, and membership in complex cellular subsystems has been shown to limit horizontal gene transfer^17,18^.

To investigate this scenario, we experimentally compared the ability of short genomic fragments to transfer resistance phenotypes towards AMPs versus antibiotics. To this end, we applied an established functional metagenomic protocol^11,19^ to identify random 1-5 kb long DNA fragments in the gut microbiome that confer resistance to an intrinsically susceptible *Escherichia coli* strain. Importantly, the length distributions of the known AMP- and antibiotic-resistance genes are well within this fragment size range (Figure S1), indicating that our protocol is suitable to capture single resistance genes for both AMPs and antibiotics. Metagenomic DNA from human gut faecal samples was isolated from two unrelated, healthy individuals who have not taken any antibiotics for at least one year. The resulting DNA samples were cut and fragments between 1-5 kb were shotgun cloned into a plasmid to express the genetic information in *Escherichia coli* K-12. About 2 million members from each library, corresponding to a total coverage of 8 Gb (the size of ~200 bacterial genomes), were then selected on solid culture medium in the presence of one of 12 diverse AMPs and 11 antibiotics at concentrations where the wild-type host strain is susceptible (Table S3). Finally, using a third-generation long-read sequencing pipeline^20^, the number of unique DNA fragments conferring resistance (i.e. resistance contigs) was determined.

In agreement with prior studies^11,21^, multiple resistant clones emerged against all tested antibiotics (Figure 2A, Table S4). In sharp contrast, no resistance was conferred against half of the AMPs tested, and, in general, the number of unique AMP resistance contigs was substantially lower than the number of unique antibiotic resistance contigs (Figure 2A, Table S4). Polymyxin B – an antimicrobial peptide used as a last-resort drug in the treatment of multidrug-resistant Gram-negative bacterial infections^22^ – is a notable exception to this trend, with a relatively high number of unique resistance contigs (Figure 2A). Indeed, a resistance gene (*mcr-1*) against Polymyxin B is rapidly spreading horizontally worldwide, representing an alarming global healthcare issue^23^. In contrast to Polymyxin B, we detected only one unique contig conferring resistance to LL37, a human AMP abundantly secreted in the intestinal epithelium^24^ (Table S4). The specific AMP resistance genes on the resistance contigs are involved in cell surface modification, peptide proteolysis and regulation of outer membrane stress response (Table 1, Table S4 and Figure S2).

**Figure 2.**
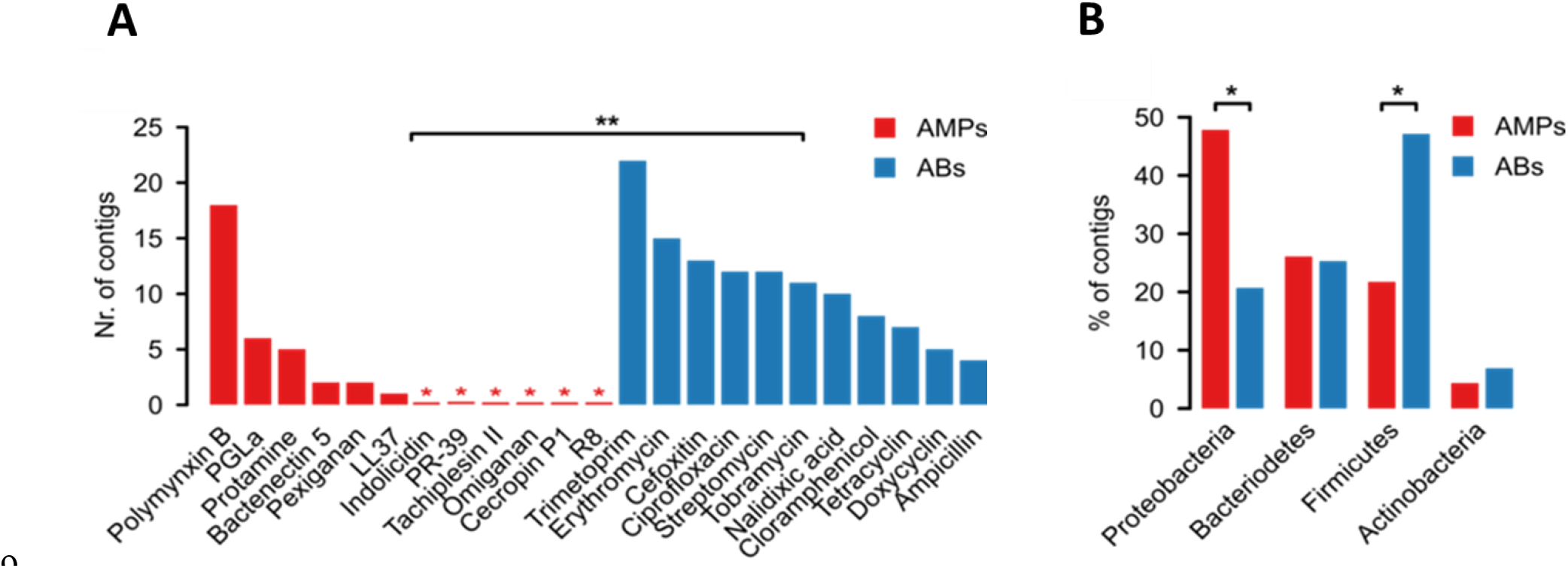
In *E. coli*, genomic fragments from the human gut microbiota confer AMP resistance less frequently than antibiotic resistance. **A)** Functional selection of metagenomic libraries with 12 AMPs (red bars) resulted in fewer distinct resistance-conferring DNA contigs than with 11 conventional small-molecule antibiotics (ABs, blue bars; P=0.002, two-sided negative binomial regression, *n*=34 (AMPs), *n*=119 (ABs)). Red asterisks indicate zero values and ** indicates a significant difference between AMPs and antibiotics. **B)** Phylum-level distribution (%) of the AMP-(red bars) and antibiotic-resistance contigs (blue bars). In the case of AMPs, significantly more resistance contigs are originating from the Proteobacteria phylum (P=0.015, two-tailed Fisher’s exact test, *n*=110), while contigs originating from phylogenetically distant relatives of the host *E. coli* from Firmicutes phylum are underrepresented (P=0.033, two-tailed Fisher’s exact test, *n*=110).

If lack of functional compatibility with the host cell prevents AMP resistance genes from exerting their phenotypic effects, then DNA fragments identified in our screen should more often come from phylogenetically closely related bacteria. 53% of the contigs showed over 95% sequence identity to bacterial genome sequences from the HMP database^25^ (see Methods, Figure S3), allowing us to infer the source taxa with high accuracy (Table S4). Indeed, AMP resistance contigs originating from Proteobacteria, which are phylogenetically close relatives of the host *E. coli*, were overrepresented (Figure 2B). Notably, this trend was not driven by polymyxin B only but was valid for the rest of the AMPs as well (Figure S4).

Whereas these patterns are consistent with the hypothesis that the genetic determinants of AMP resistance are difficult to transfer via short genomic fragments owing to a lack of functional compatibility with the new host, another explanation is also plausible. In particular, AMP resistant bacteria might be relatively rare in the human gut microbiota, therefore, AMP resistance genes from these bacteria might simply remain undetected. However, as explained below, we can rule out this alternative hypothesis.

### AMP-resistant gut bacteria are abundant and phylogenetically diverse

To assess the diversity and the taxonomic composition of gut bacteria displaying resistance to AMPs and antibiotics, we carried out anaerobic cultivations and selections of the gut microbiota using a state-of-the-art protocol^26^. To this end, faecal samples were collected from 7 healthy individuals (i.e. Fecal 7 mix, see Methods). As expected^26^, the cultivation protocol allowed representative sampling of the gut microbiota: we could cultivate 65-74% of the gut microbial community at the family level in the absence of any drug treatment (Figure S5, Table S5). Next, the same faecal samples were cultivated in the presence of one of 5-5 representative AMPs and antibiotics, respectively (Table S6). We applied drug dosages that retained 0.01 to 0.1% of the total cell populations from untreated cultivations (Table S6) and assessed the taxonomic composition of these cultures by 16S rRNA sequencing (see Methods).

Remarkably, the diversities of the AMP-treated and the untreated bacterial cultures did not differ significantly from each other (Figure 3A), despite marked differences in their taxonomic compositions (Figure 3B). AMP-treated samples contained several bacterial families from the Firmicutes and Actinobacteria phyla, which are phylogenetically distant from *E. coli* (Figure 3C). Notably, exposure to AMP stress provided a competitive growth advantage to bacterial families that remained undetected in the untreated samples (Figure 3C). The examples include Desulfovibrionaceae, *a* clinically relevant family that is linked to ulcerative colitis^27^ – an inflammatory condition with elevated AMP levels^28^ (Figure 3C). In sharp contrast, the diversity of antibiotic-treated cultures dropped significantly compared to both the untreated and the AMP-treated cultures (Figure 3A and Figure S6). Several bacterial families had a significantly lower abundance in the antibiotic-treated cultures than in the untreated ones (Figure 3C). These results indicate that the human gut is inhabited by taxonomically diverse bacteria that exhibit intrinsic resistance to AMPs.

**Figure 3.**
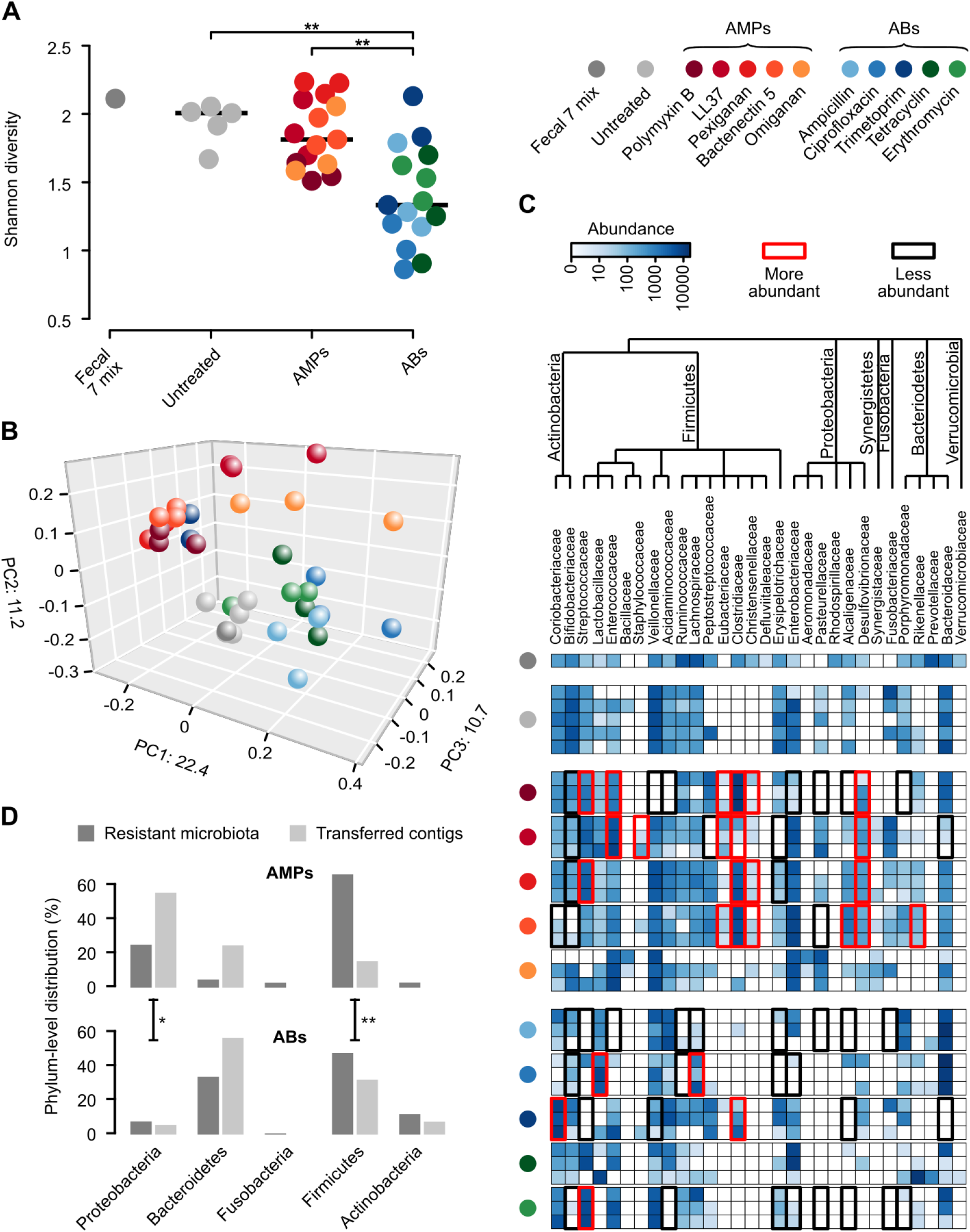
Culturing reveals a diverse AMP-resistant gut microbiota with limited potential to transfer resistance to *E. coli*. **A)** Diversities of the cultured microbiota and the original faecal sample (Fecal 7 mix). Data represents Shannon alpha diversity indexes at family level based on 16S rRNA profiling of V4 region. AMP/antibiotic treatments are colour-coded. Untreated samples were grown in the absence of any AMP or antibiotics. ** indicate a significant difference, P<0.01 from two-sided Mann-Whitney U test, *n*=36. Central horizontal bars represent median values. **B)** A PCoA (principal coordinate analysis) plot based on unweighted UniFrac distances^33^ separates the AMP- and antibiotic-resistant, and the untreated microbial communities (P=0.001, PERMANOVA test, *n*=36). **C)** Differential abundance analyses between the untreated and the AMP/antibiotic-resistant microbiota at the family level (see Methods). Brackets depict a significant increase (red) or decrease (black) in abundance of a given family as a consequence of AMP or antibiotic treatment. **D)** Phylum-level distributions of resistant gut bacteria and resistance DNA contigs originating from them. Compared to their relative frequencies in the drug-treated cultured microbiota, the phylogenetically close Proteobacteria contributed disproportionally more AMP than AB resistance contigs, whereas the opposite pattern was seen for the distantly related Firmicutes. Asterisks indicate significant interaction terms in logistic regression models, P<0.05 (*) and P<0.01 (**), respectively (for more details see Methods).

### Human microbiota harbours a large reservoir of AMP resistance genes

Next, we assessed if the high taxonomic diversity in the AMP-resistant microbiota corresponds to a diverse reservoir of AMP resistance genes. To this end, we annotated previously identified AMP- and antibiotic-resistance genes in a set of gut bacterial genomes^15^ representing bacterial families that were detected in our culturing experiments upon AMP- and antibiotic-selection, respectively (Figure 3C, for details, see Methods). Remarkably, 65% of our literature curated AMP resistance gene families (Table S1) were represented in at least one of these genomes (Table S2), which is similar to that of antibiotic resistance gene families (58%). Finally, AMP resistance gene families, on average, were 32% more widespread in these species than the same figure for antibiotic resistance genes (Figure S7). Thus, the human gut harbours diverse AMP-resistant bacteria and a large reservoir of AMP resistance genes.

### Phylogenetic constraints on the functional compatibility of AMP resistance genes

We next directly tested whether the shortage of AMP resistance DNA fragments from distantly related bacteria can be explained by the low potential of genomic fragments to transfer AMP resistance phenotypes to *E. coli*. To this end, we constructed metagenomic libraries from the AMP- and antibiotic-resistant microbiota cultures. From each AMP and antibiotic treatments, two biological replicates were generated (see Methods), resulting in 10-10 libraries, covering 25.6 Gb and 14 Gb DNA, respectively (Table S7). These metagenomic libraries were next screened on the corresponding AMP- or antibiotic-containing solid medium. Finally, the phylogenetic sources of the resulting AMP- and antibiotic-resistance contigs were inferred (Figure S8, Table S8). Compared to their relative frequencies in the drug-treated cultured microbiota, the phylogenetically close Proteobacteria contributed disproportionally more AMP- than antibiotic-resistance DNA fragments, whereas the opposite pattern was seen for the distantly related Firmicutes (Figure 3D, Table S9). Taken together, phylogenetically diverse gut bacterial species show AMP resistance, but there is a shortage of transferable AMP resistance DNA fragments from phylogenetically distant relatives of *E. coli*.

### Pervasive genetic background dependence of AMP resistance genes

Finally, we present evidence that DNA fragments that confer resistance to AMPs and were isolated from our screens show stronger genetic background dependence than those conferring resistance to antibiotics.

To test the generality of genetic background-dependency of AMP resistance genes, we examined how DNA fragments that provide AMP- or antibiotic-resistance in *E. coli* influence drug susceptibility in a related Enterobacter species, *Salmonella enterica*. We analyzed a representative set of 41 resistance DNA fragments derived from our screens (Table S10) by measuring the levels of resistance provided by them in both *E. coli* and S. *enterica*. Strikingly, while 88% of the antibiotic resistance DNA fragments provided resistance in both host species, only 38.9% of AMP resistance DNA fragments did so (Figure 4A, Table S10). Thus, the phenotypic effect of AMP resistance genes frequently depends on the genetic background, even when closely related hosts are compared.

**Figure 4.**
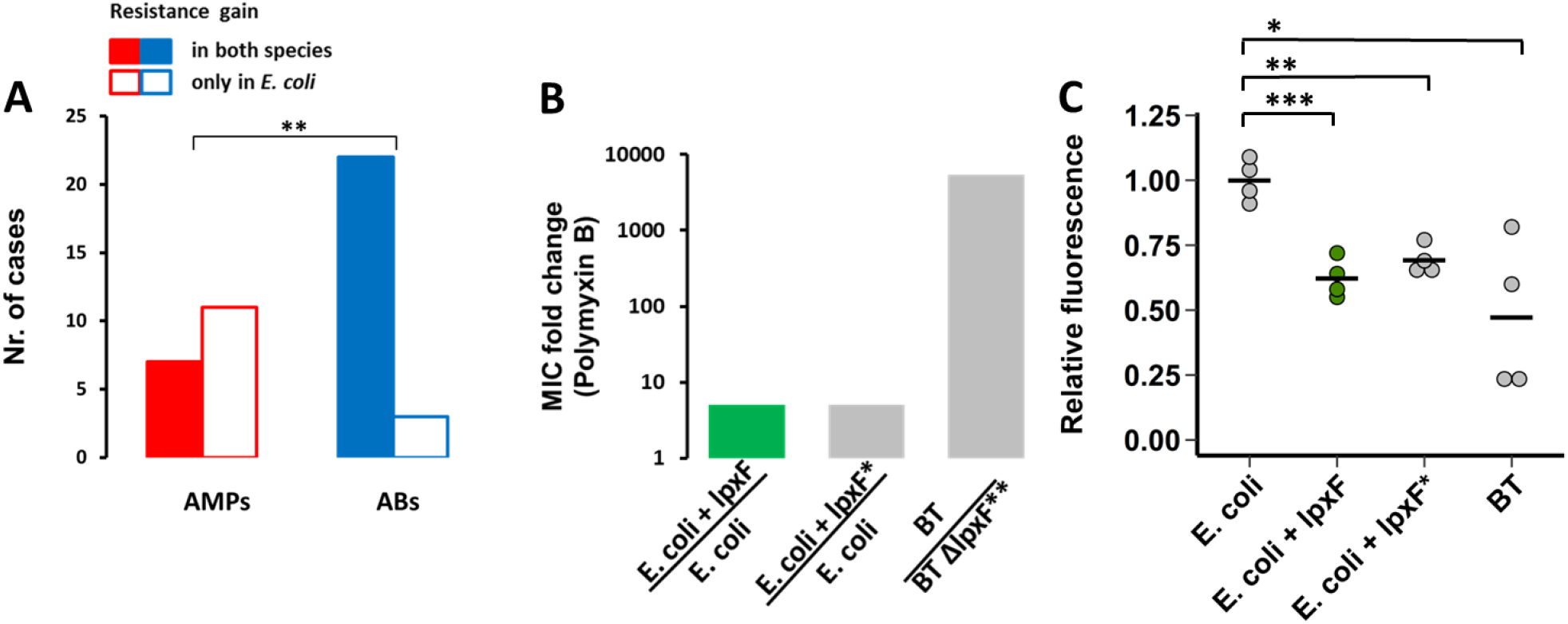
AMP resistance DNA fragments provide host-dependent phenotypic effects. **A)** A significantly lower proportion of AMP resistance DNA fragments conferred resistance in both *E. coli* and S. *enterica* compared to antibiotic resistance DNA fragments, suggesting weaker between-species conservation of the AMP resistance phenotypes. Asterisks indicate significant difference, P=0.0011, two-sided Fisher’s exact test, *n*=16 for AMPs, *n*=25 for ABs. **B)** The *lpxF* ortholog from *Parabacteroides merdae* (marked as lpxF) isolated in our screen (Table 1) increases Polymyxin B resistance of *E. coli* only five-fold (green bar), to the same extent as a previously characterized fully functional *LpxF* from *Francisella tularensis* (marked as *lpxF*\*). In contrast, lpxF in its original host, *B. thetaiotaomicron* (denoted as *lpxF*\*\*) provides a 5000-fold increment in polymyxin B resistance. **C)** The *lpxF* and *lpxF*\* decrease the net negative surface charge of *E. coli* (green bar) to the same extent, to the level of wild-type *Bacteroides thetaiotaomicron* (BT) expressing its native *LpxF*\*\*. The fluorescence signal is proportional to the binding of the FITC-labeled poly-L-lysine polycationic molecule. Less poly-L-lysine binding reflects a less negative net cell surface charge. *, **, *** indicate significant differences (P= 0.03, 0.001 and 0.0004, respectively, Welch Two-Sample t-test, n=4 biological replicates, central horizontal bars represent mean values). Corresponding microscopic pictures are shown in Figure S9.

**Table 1.** List of putative AMP resistance genes identified from our functional metagenomic screens. These resistance genes could be functionally annotated based on a literature-curated catalogue of resistance genes (Table S1).

| Resistance gene | Resistance gene function | Classification of resistance function | Estimated source organism | UniProt ID | AMP |
| --- | --- | --- | --- | --- | --- |
| *lpxF* | Lipid A 4'-phosphatase | Target alteration | Parabacteroides sp. | P94571 | Polymyxin B, Pexiganan |
| *lpxF* | Lipid A 4'-phosphatase |  | NA | P75806 | Polymyxin B |
| *lpxF* | Lipid A 4'-phosphatase |  | Parabacteroides merdae ATCC 43184 | NA | Polymyxin B, Pexiganan |
| *arnTEF* | 4-amino-4-deoxy-L-arabinose-modification of Lipid-A |  | Escherichia coli MS 107-1 | P76473<br>Q47377<br>P76474 | Polymyxin B |
| *pmrAB (qseBC)* | Response regulator | Regulation | Sutterella wadsworthensis 2_1_59BFAA | P52076<br>P40719 | Polymyxin B |
| *pmrD* | Signal transduction protein |  | Escherichia coli MS 107-1 | P37590 | Polymyxin B |
| *ompT* | Protease 7 precursor | Agent inactivation | Escherichia coli MS 196-1 | P09169 | Bactenectin 5, LL37, Pexiganan |

As an example, we finally focused on a putative ortholog of a previously characterized AMP resistance gene, *lpxF*^12^. LpxF is a key determinant of AMP resistance in Bacteroidetes, a member of the human gut microbiota. By decreasing the net negative surface charge of the bacterial cell, it provides a 5000-fold increment in Polymyxin B resistance in these species^12,29^. To test the genetic background dependence of this resistance gene, we expressed the *lpxF* ortholog in *E. coli* and we found that it provides a mere five-fold increase in Polymyxin B resistance (Figure 4B). Reassuringly, surface charge measurements proved that this *lpxF* is fully functional in *E. coli* (Figure 4C, Figure S9). The compromised resistance phenotype conferred by *lpxF* in the new host shows that the function of other genes is also required to achieve the high AMP resistance level seen in the original host.

## Discussion

This work systematically investigated the mobility of AMP versus antibiotic resistance genes in the gut microbiome. We report that AMP resistance genes are less frequently transferred between members of the gut microbiota than antibiotic resistance genes (Figure 1). In principle, this pattern could be explained by at least two independent factors: shortage of relevant selection regimes during the recent evolutionary history of the gut microbiota and lack of functional compatibility of AMP resistance genes upon transfer to a new host. We focused on testing the second possibility due to its experimental tractability and relevance to forecast the mobility of resistance genes upon AMP treatment. In a series of experiments, we showed that phylogenetically diverse gut bacteria display high levels of AMP resistance (Figure 3), yet the underlying resistance genes often fail to confer resistance upon transfer to *E. coli* (Figure 2 and Figure 3). Furthermore, we demonstrated that the AMP resistance conferred by genomic fragments often depends on the genetic background of the recipient bacterium (Figure 4). Together, these results support the notion that horizontal acquisition of AMP resistance is constrained by phylogenetic barriers owing to functional incompatibility with the new host cell^30^. We speculate that the large differences in functional compatibility between antibiotic- and AMP-resistance genes might be caused by the latter being more often parts of highly interconnected cellular subsystems, such as cell envelope biosynthesis pathways. Clearly, deciphering the biochemical underpinning of functional incompatibility remains an area for future research.

We note that compromised benefit may not be the only manifestation of functional incompatibility and the exclusive reason for the limited presence of AMP resistance genes in the mobile gene pool (Figure 1). It is also plausible that some AMP resistance genes have severe deleterious side effects in the new host in addition to conferring a compromised resistance. For example, the introduction of *lpxF* into bacterial pathogens reduces virulence in mice, probably because it perturbs the stability of the bacterial outer membrane in enterobacterial species^31^. Indeed, LpxF increases the sensitivity of *E.coli* to bile acid, a membrane-damaging agent secreted into the gut of vertebrates (Figure S10). Future works should elucidate whether AMP resistance genes are especially prone to induce deleterious side effects compared to antibiotic resistance ones.

An important and unresolved issue is why natural AMPs that are part of the human innate immune system have remained effective for millions of years without detectable resistance in several bacterial species. One possibility, supported by our work, is that the acquisition of resistance through horizontal gene transfer from human gut bacteria is limited, most likely due to compromised functional compatibility in the recipient bacteria. We do not wish to claim, however, that AMPs in clinical use would generally be resistance-free. In agreement with the prevalence of Polymyxin B resistance DNA fragments (Figure 2A), a plasmid conferring colistin resistance is spreading globally^32^. Rather, our work highlights major differences in the frequencies and the mechanisms of resistance across AMPs, with the ultimate aim to identify antimicrobial agents less prone to resistance. Finally, our results highlight a previously ignored potential problem with the clinical usage of AMPs. Our work indicates that upon various AMP stresses, the abundance of bacterial families traditionally associated with inflammatory bowel diseases (e.g. Desulfovibrionaceae) increases (Figure 3C). Future works should examine whether AMP stress increases the risk of human diseases by specifically perturbing the human microbiota composition.

## Acknowledgements

We thank Dezsö Módos, Dávid Fazekas, József Sóki and Edit Urbán for their technical support. Funding: This work was supported by the ‘Lendület’ programme of the Hungarian Academy of Sciences (B.P. and C.P.), the Wellcome Trust (B.P.), H2020-ERC-2014-CoG (C.P.), GINOP-2.3.2-15-2016-00014 (EVOMER, C.P. and B.P.) and GINOP-2.3.2-15-2016-00020 (MolMedEx TUMORDNS, C.P.), NKFIH grant K120220 (B.K.), NKFIH grant FK124254 (O.M.). BK holds a Bolyai Janos Scholarship.

## Data and materials availability

All data is available in the main text or the supplementary materials.

## Materials and Methods

### Establishing a comprehensive AMP resistance gene dataset

Even though several databases have been created for antibiotic resistance genes, a comprehensive list of AMP resistance genes has not been compiled so far. Therefore, we carried out a systematic literature mining in PubMed NCBI and Google Scholar with the keywords “antimicrobial peptide” + “resistance”. From the identified publications genes with experimentally confirmed influence on AMP susceptibility were included in our manually curated AMP resistance gene dataset (Table S1). Altogether 131 AMP resistance genes were identified. As a next step, the compiled AMP resistance genes were classified into resistance gene families (orthologous gene groups or orthogroups) by the *EggNOG-mapper* software (version 0.12.7) with the bacterial EggNOG 4.5.1 database^34^. Then, AMP resistance genes were classified into broad functional categories analogous to the classification of antibiotic resistance genes in The Comprehensive Antibiotic Resistance Database (CARD)^35^. To obtain a comparable dataset for known antibiotic resistance genes we downloaded the CARD database^35^. Antibiotic resistance genes from CARD were grouped into resistance gene families in the same way as AMP resistance genes using the EggNOG 4.5.1 database (Table S1).

### Analysis of the mobile gene pool of the gut microbiota

A previously published mobile gene catalogue of the human gut microbiota^15^ was analyzed to compare the patterns of horizontal gene transfer events involving AMP and antibiotic resistance genes across a wide range of bacteria. This mobile gene catalogue relies on the identification of nearly identical genes in distantly related bacterial genomes and thereby provides a snapshot on the gene set subjected to recent horizontal gene transfer events in a representative sample of the gut microbiota. The goal in our analysis was to determine the presence/absence pattern of the AMP and antibiotic resistance genes not only in the mobile gene pool but also in the 567 genomes from which the mobile gene pool was derived. In this way, not only the horizontally transferred resistance genes were identified but also those that have not been detected in such transfer events, but were present in the gut microbiome. To this end, the genomes and proteomes used by Brito et al.^15^ were downloaded from the Human Microbiome Project (HMP) database (https://www.hmpdacc.org/HMRGD/) and from Fijicomp website (http://www.fijicomp.org). DNA sequences derived from the latter database were used for ORF prediction with *Prodigal* software (version 2.6.3,^36^). Then, a sequence similarity search was applied on the compiled list of proteins encoded in the analyzed genomes to identify those that were present in the mobile gene pool as well. The sequence similarity search between the mobile gene pool and the proteins from the genomes was carried out with the *blastx* option of the *Diamond* software (version 0.9.10,^37^) with 50% sequence coverage and 100% sequence identity (parameters were chosen to reproduce the original publication of the mobile gene pool^15^). Out of the 37,870 unique mobile genes in the mobile gene pool, we identified 37,184 in the genomes (98.28%).

Next, both the antibiotic- and AMP-resistance genes were identified among the mobile genes and among those that have not been detected in the mobile gene pool but were present in the genomes. For this functional annotation, a blast search was carried out against the antibiotic resistance genes from the CARD database^35^ with the *blastp* option of the *Diamond* software^37^ with strict parameters (*e*-value < 10^-5^, > 40% identity at the protein level and 80% query sequence coverage) (Table S1). In a similar vein, AMP resistance genes were identified by performing *blastp* sequence similarity search against the manually curated list of AMP resistance genes (Table S1). This AMP resistance gene list was compiled by literature mining in PubMed NCBI and Google Scholar with the keywords “antimicrobial peptide” + “resistance”. From these publications genes with experimentally confirmed influence on AMP susceptibility were included in our AMP resistance gene database (Table S1). Antibiotic and AMP resistance genes in our databases were classified into resistance gene families by the *EggNOG-mapper* software (version 0.12.7) on the bacterial EggNOG 4.5.1 database^34^ (Table S1). For the annotated resistance gene list in the mobile gene pool, see Table S2.

To compare the relative frequency of the AMP- and antibiotic-resistance genes in the mobile gene pool (Figure 1A), we restricted the analysis to one genome per species. This was necessary to avoid sampling bias since different species were represented by unequal numbers of genomes in the dataset. To this aim, 16S rRNA sequences were determined for each genome (for HMP genomes they were downloaded from the Silva database^38^, while for Fiji genomes they were identified directly in the genomes using the *RNAmmer* software (version 1.2 ^39^). Then, genomes with lower than 2% 16S RNA gene dissimilarities were collapsed into genome groups (‘species’ or OTUs) using average linkage clustering as it is described in the publication of the mobile gene pool^15^. Then, from each genome group, the genome with the largest number of annotated resistance genes was selected for further analyses. Resistance genes in the mobile gene pool that resulted in a blast hit both from the AMP- and the antibiotic-resistance databases were excluded from the analysis. The remaining annotated resistance genes were counted and plotted in Figure 1A.

For each unique mobile AMP- or antibiotic-resistance gene we estimated the minimum number of independent transfer events (Figure 1B) by counting the number of genome groups (i.e. OTUs) in which the gene is present in the mobile gene pool^15^.

### Construction of gut metagenomic libraries

To sample the gut resistome, we applied a previously established small-insert shotgun metagenomic protocol^11^ with small modifications. This method identifies small genomic fragments that decrease drug susceptibility via plasmid-mediated gene transfer. For the construction of the metagenomic libraries, human stool samples were obtained from two healthy unrelated individuals, who have not taken any antibiotics for at least one year prior to sample donation. Samples were handled with the observation of ethical rules. Gut community DNA was isolated immediately after sample donation, using ZR Fecal DNA MiniPrep™ kit (Zymo Research) according to the manufacturer’s instructions (http://www.zymoresearch.com/downloads/dl/file/id/91/d6010i.pdf). Subsequently, 10 μg of metagenomic DNA from each sample was partially digested with 0.25 U MluCI restriction enzyme (New England BioLabs) in 10x CutSmart Buffer (New England BioLabs) at 37°C for 20 minutes, followed by heat inactivation at 85°C for 20 min. MluCI is a four-base cutter restriction enzyme that produces overhangs complementary to the ones that EcoRI produces. By varying incubation time or the enzyme concentration, the size range of the resulting DNA fragments can be set. The fragmented DNA was size selected by electrophoresis on a 1 % (m/V) agarose gel in 1X Tris-Acetate-EDTA (TAE) buffer. A gel slice corresponding to 1500-5000 bp was excised from the gel and DNA was isolated using a GeneJET Gel Extraction and DNA Cleanup Micro Kit (Thermo Scientific). 5 μg of pZErO-2 plasmid DNA (Thermofisher, https://tools.thermofisher.com/content/sfs/vectors/pzero2_map.pdf) was digested with 25 U EcoRI restriction enzyme (Fermentas) in 10x EcoRI Buffer (Fermentas) for 2 hrs, followed by 20 minutes heat inactivation at 65°C. After purification with DNA Clean & Concentrator™-5 kit (Zymo Research), digested pZErO-2 plasmid was dephosphorylated with FastAP alkaline phosphatase (Thermo Scientific) as follows: 4 μg plasmid DNA was incubated with 4U enzyme in 10x FastDigest buffer at 37°C for 1 hr, followed by 5 minutes heat inactivation at 74°C and purification with DNA Clean & Concentrator™-5 kit (Zymo Research). DNA was ligated into pZErO-2 at the EcoRI site using the Rapid DNA ligation kit (Thermo Scientific). The ligation reaction was performed in 15 μl total volume using a 5:1 insert-vector ratio: 4.5 μl (310 ng) digested and gel purified DNA insert, 0.65 μl (62 ng) EcoRI-cut pZErO-2 vector, 3 μL 5X ligation buffer, 0.75 μL 10 mM ATP, 4.1 μL dH_2_O, 2 μL T4 DNA Ligase (5 U/μl). The ligation mixture was incubated at 16°C overnight, followed by heat inactivation at 65°C for 10 minutes.

Prior transformation, the ligation mixture was purified with DNA Clean & Concentrator™-5 kit (Zymo Research). 3.5 μL of the resulting ligation mixture was transformed by electroporation into 50 μL of electrocompetent *E. coli* DH10B™ cells (Invitrogen). Electroporation was carried out with a standard protocol for a 1 mm electroporation cuvette. Cells were recovered in 1 mL SOC medium, followed by 1-hour incubation at 37°C. 500 μL of the recovered cells was plated onto square Petri dishes containing Luria Bertani (LB) agar supplemented with 50 μg/mL kanamycin. In order to assess the library size (number of colony forming units (CFU)), 1 μL of the electroporated cells was saved for plating onto a separate Petri dish containing Luria Bertani (LB) agar supplemented with 50 μg/mL kanamycin. From each plate, 10 clones were randomly picked for colony PCR in order to confirm the presence and the size distribution of the inserts. PCR was performed using the Sp6-T7 primer-pair (Table S11) flanking the EcoRI site of the multiple cloning site of the pZErO-2 vector. The sizes of the PCR products were determined by gel electrophoresis and the average insert size was calculated as 2-3 kb. The size of each library was determined by multiplying the average insert size by the number of total colony forming units (CFU). The size distributions of the libraries varied between 4.4 – 16 Gb coverage with this protocol, which is in line with a previously published state-of-the-art protocol ^11,19^. The resulting colonies from the Petri dishes were collected and the plasmid library was isolated using InnuPREP Plasmid Mini Kit (Analytic Jena). 30-60 ng of isolated plasmid library was transformed by electroporation into 40 μL electrocompetent *E. coli* BW25113 (prepared as described in ^40^). This *E. coli* strain was used for the functional selections (see below). After electroporation, cells were recovered in 1 mL of LB medium for 1 hour at 37°C. Special care was taken to achieve high electroporation efficiency in order to cover 10-100 times the original library size. In this way, we ensured that most library members are electroporated from the plasmid library. The 1 mL recovered cell culture was added to 9 ml of LB medium supplemented with 50 μg/mL kanamycin, and grown at 37 °C for 2-3 hours until it reached the 7.5-10 x 10^8^ cell density (OD_600_: 1.5-2). Cell aliquots were frozen in 20% glycerol and kept at −80°C for subsequent functional selection experiments.

Metagenomic libraries were generated from the uncultured microbiota (i.e. the total DNA extracted from the stool samples) and from the cultured microbiota (i.e. the genomic DNA extracted from the cultured pooled microbiota), too. For details see the section *“Cultivation of the gut microbiota under anaerobic conditions and DNA extraction*”.

### Functional metagenomic selections for AMP/AB resistance

Functional selections for resistance were carried out on solid plates containing one of the 12 antimicrobial peptides (AMPs) or 11 antibiotics (ABs) (Table S3). Instead of the plating assay that is commonly used in the field^11^, we applied a modified gradient plate assay^41^, where bacteria are exposed to a concentration gradient of the antimicrobial instead of a single concentration. We found that this strategy improves reproducibility of AMP selections, where changes in the resistance levels are relatively small compared to that in the case of ABs. The growth medium in these plates was a modified minimal salt medium (MS) with reduced salt concentration (1 g (NH4)_2_SO_4_, 3 g KH_2_PO_4_, 7 g K_2_HPO_4_, 100 μl MgSO_4_ (1 M), 540 μl FeCl_3_ (1 mg/ml), 20 μl thiamine (50 mg/ml), 20 ml casamino acids (BD) (10 % (m/V)), 5 ml glucose (40 % (m/V)) in a final volume of 1 L), since most AMPs are not effective *in vitro* in the presence of high salt concentrations. In the case of the AMP containing plates, the solidifying agent was changed to 1.5 % (m/V) of low melting point agarose (UltraPure™ LMP Agarose (Invitrogen)) from 1.5 % (m/V) agar, in order to prevent any heat-induced structural damage of the peptides during plate pouring. ABs and AMPs were purchased from Sigma and ProteoGenix, respectively. Onto each of the gradient plates (Tray plates, SPL Life Sciences) 2×10^8^ cells were plated out from the thawed stocks of *E. coli* BW25113 bearing the metagenomic plasmid libraries. In this way, each metagenomic library member was represented about 10-100 times on each plate. We found this necessary for a good reproducibility for our experiments. Subsequently, plates were incubated at 30°C for 24 hours. For each functional selection, a control plate was prepared where the same number of *E. coli* BW25113 was plated out. These cells contained the pZErO-2 plasmid with a random metagenomic DNA insert that has no effect on AMP/AB resistance. This control plate showed the minimum inhibitory concentration (MIC) of the antimicrobial without the effect of a resistance plasmid. The empty plasmid was not applicable as a control because in the absence of a DNA insert the CcdB toxic protein is expressed from the plasmid. In order to isolate the resistant clones from the library plates, sporadic colonies were identified above the MIC level (defined using the control plate) by visual inspection. These clones were collected by scraping them into 2 ml of LB broth and stored subsequently at −80°C.

### Validation of the resistance-conferring metagenomic DNA fragments

Following selection of the metagenomic libraries, the putative resistance phenotypes conferred by the plasmid selections were confirmed for a representative fraction of the colonies. From each selection at least 20 colonies were picked and the MIC increase was determined by a standard broth microdilution method^42^, as it is described in the section “*Quantification of the resistance gains that metagenomic contigs provide*”. For these measurements, the same control plasmid was used as in the functional selections. MIC values were determined after 16-24 hours of incubation at 30°C with a continuous shaking at 240 rpm. Plasmids from validated resistant clones were retransformed into the BW25113 *E. coli* strain, followed by a second MIC determination in order to exclude the possibility that the increase in the MIC was induced by genomic mutations. Plasmids not showing MIC increase in the validation protocol were excluded from further analysis. Mostly the clones situated closer than 1 cm to the MIC level on the gradient plates did not confer resistance during validation. To avoid these false positive resistance plasmids, colonies at the borderline were not collected for further analysis without confirming their phenotype. The rest of the colonies were collected by scraping them into 2 ml of LB broth. Bacterial samples then were stored at −80°C in 20 % (m/V) glycerol. When only a few clones were on the plates, all were tested for resistance to make sure that we do not lose potential hits. We encountered such situations only in the case of AMPs. If the number of resistant clones on a plate was less than or equal to 20, plasmids were isolated individually from the MIC validated clones and sent for Sanger bidirectional sequencing with the Sp6-T7 primer pair (Table S11). When the resistance-conferring insert was longer than what the initial sequencing covered, we applied a primer walking strategy to sequence the middle part of the insert.

### Amplification of the resistance-conferring metagenomic DNA fragments

Plasmid pools from the scraped resistant clones were PCR amplified for subsequent PacBio sequencing^43^. To this aim, first, the plasmid pools were isolated from each metagenomic selection using InnuPREP Plasmid Mini Kit (Analytic Jena). Then, these plasmid pools served as templates for subsequent PCR amplification reactions to amplify the inserts from the pooled plasmids. These PCR reactions were performed with barcoded Sp6 and T7 specific primers as forward and reverse primers, respectively. The 16-base long PacBio barcode dual-end labelling scheme was used to label each plasmid pool for sample identification in the subsequent PacBio sequencing. Primer sequences are shown in Table S11. PCR reactions consisted of 30 ng of template DNA, Q5 Hot Start High-Fidelity 2x Master Mix (New England BioLabs), 0.2 μM barcoded primers in a final reaction volume of 25 μl. Following an optimization process, the number of PCR cycles was reduced to 15 in order to minimize amplification bias. The following thermocycler conditions were used: 98°C for 5 minutes, 15 cycles of 98°C for 15 seconds + 69°C for 30 seconds + 72°C for 2 minutes, and 72°C for 7 minutes. The amplified metagenomic inserts were then cleaned using the DNA Clean & Concentrator™-5 kit (Zymo Research) and their concentration was measured with Qubit fluorometer (Invitrogen). In order to get rid of the short amplicons (e.g. primer dimers), which may interfere with the sequencing process, barcoded amplicons were mixed at an equimolar ratio and the sample was gel extracted following electrophoreses using 1% agarose gel. The sample was purified (Zymoclean™ Gel DNA Recovery Kit) prior sequencing.

### Pacbio sequencing and data analysis

Sequences of the pooled PCR products were obtained from the Norwegian Sequencing Centre at the University of Oslo, Norway. The library was prepared using Pacific Biosciences Amplicon library preparation protocol. Samples were sequenced with Pacific Biosciences RS II instrument using P6-C4 chemistry and MagBead loading in one SMRT cell. 61,641 reads were obtained with a mean length of 20,961 bp. Reads were filtered without demultiplexing using *RS_subreads.1* pipeline on *SMRT Portal* (version 2.3.0) using default settings (number of passes 1, minimum accuracy 0.9). Following barcode detection and demultiplexing, reads were collapsed to consensus sequences using the long amplicon analysis pipeline of the *SMRT Portal* with default settings. We validated our sequencing effort on a mock sample containing 9 previously sequenced DNA contigs that originate from our metagenomic selections. Reassuringly, out of the 9 sequences 8 were present in the Pacbio sequencing result with at least 99% sequence identity. The single non-detected contig was the longest one (4500 bp) which may indicate a bias of the pipeline toward shorter insert sizes.

### Functional annotation and resistance gene identification on the metagenomic contigs

To functionally analyze the ORFs on the assembled contigs from the metagenomic selections, ORFs were predicted and annotated using the *Prokka* suite (version 1.11, ^44^) with the bacterial prediction settings and an *e*-value threshold of 10^-5^. Within *Prokka, Prodigal* (version 2.6.3, ^36^) subscript was modified to run without -*c* parameter to identify highly probable ORFs, even if the ends were not closed. This was necessary because some contigs may have been shortened by a few residues during the cloning process or in the assembly due to low coverage, without losing their functionality. Other parameters were kept as default. Next, ORFs on the metagenomic contigs were functionally annotated with our resistance gene lists introduced in the section “*Analysis of the mobile gene pool of the gut microbiota*”. Specifically, an ORF was classified as an antibiotic resistance gene when sequence similarity search using *blastp*^45^ against The Comprehensive Antibiotic Resistance Database (CARD)^35^, resulted in an annotation with an *e*-value < 10^−5^, identity > 30% and coverage > 80%. Here, the minimum sequence identity was lower than in the case of the analysis of the mobile gene pool, since the experimentally observed resistance phenotype provided an extra confidence for the annotation. Similarly, AMP resistance genes were identified by performing *blastp* sequence similarity search against the manually curated list of AMP resistance genes (Table S1).

To estimate the identity of the donor organisms from which the assembled DNA contig sequences originated from, a nucleotide sequence similarity search was carried out for the entire DNA contigs as query sequences against the genome sequences from the Human Microbiome Project^25^ using *blastn*, with an *e*-value threshold of <10^-10^. Taxonomy was assigned with the *ete3* toolkit^34^.

### Cultivation of the gut microbiota under anaerobic conditions and DNA extraction

In order to compare the abundance and phylogenetic distribution of the AMP and AB resistant gut bacteria, we applied a recently established anaerobic cultivation protocol of the human gut microbiota with small modifications^26^. For this purpose, human faecal samples were obtained from seven healthy unrelated volunteers, who had not taken any antibiotics for at least one year prior to sample donation. Ethical rules were observed throughout the whole study. Following defecation, stool samples were immediately placed into uncapped 50 ml plastic tubes (Sarstedt), deposited in anaerobic bags (Oxoid, Thermo Scientific) and samples were transferred into the anaerobic chamber within 1 hour after sample collection. All anaerobic experiments were performed in a Bactron IV anaerobic chamber (Shel Lab) filled with an atmosphere of 95% nitrogen and 5% hydrogen, with palladium catalysts. Two grams of the faecal samples were suspended in 20 ml of modified Gifu Anaerobic Medium (GAM) Broth (HyServe). After 10 minutes of incubation, letting the solid particles settle down, the supernatants were supplemented with 20% glycerol, aliquoted and stored at −80°C. Prior to the cultivation of the microbiota, thawed aliquots from the samples of the 7 individuals were combined in an equal ratio (we refer to this sample mix as “Fecal 7 mix” sample) in the anaerobic chamber. Then this Fecal 7 mix sample was plated out anaerobically in the presence and absence of one of the five AMPs or one of the five ABs that were active in the culturing medium (Table S3). The culturing medium was modified Gifu Anaerobic Medium (GAM), considering that the best reconstruction of the composition and architecture of the human gut bacterial community could be obtained using this medium^26^. In the case of AMP containing plates, the solidifying agent was low melting agarose instead of agar for the same reason as before (see section “*Functional metagenomic selections for AMP/AB resistance”*). Each AMP/AB was applied in three different concentrations on separate plates in three replicates. Plates were incubated at 37°C in the anaerobic chamber for a 4-day time interval, since following the 4^th^ day we observed a plateau in the number of appearing colonies. Following colony counts the plates had to fulfil two requirements to be selected for subsequent analysis. First, colony numbers needed to be in the range of 0.01-0.1% as compared to the colony numbers in the absence of any AMP/AB treatment (we refer to these samples as Untreated). Second, colony numbers needed to be high, but still in the countable range (1000-5000 colonies). To be in this range in the case of the untreated samples as well the Fecal 7 mix sample was plated out in a concentration that is 1000-fold more dilute as what was applied in the case of AMP/AB selections. For Trimetoprim the selection pressure could not be increased that much to select for the most resistant 0.01-0.1% of the population since 0.49 % of the colonies were able to grow even at the solubility limit of this antibiotic. From the selected plates resistant colonies were scraped and the pooled colonies from each plate were used to isolate genomic DNA with ZR Fecal DNA MiniPrep™ kit (Zymo Research) according to the manufacturer’s instructions (http://www.zymoresearch.com/downloads/dl/file/id/91/d6010i.pdf). The isolated DNA samples were subsequently used for both small-insert shotgun library constructions (referred to them as libraries from cultured microbiota, Table S7) and for 16S rRNA based community profiling. To estimate the relative resistance level of the gut microbiota compared to *E. coli* BW25113, we used the colony counts from the anaerobic cultivation experiments at each AMP/AB concentration, i.e. the resistance level of the gut microbiota against an AMP/AB is defined as the AMP/AB concentration at which only 0.1-0.01% of the untreated microbiota population could grow (see details above), analogously to an MIC value determination for a single species. The susceptibility of *E. coli* was estimated in the same way as for the gut microbiota by plating out anaerobically the same number of cells as in the case of the Fecal 7 mix sample for each AMP/AB treatment. Then we determined the AMP/AB concentrations at which only 0.1-0.01% of the *E. coli* cells could grow. The relative resistance level of the gut microbiota was defined as the resistance level of the gut microbiota divided by the resistance level of *E. coli* (Table S6).

### 16S rRNA-based bacterial community profiling

To determine the taxonomic composition of the AMP/AB resistant gut bacterial communities, we sequenced and analyzed the V4 region of the 16S rRNA genes from the Fecal 7 mix samples cultivated in the presence of individual AMPs and ABs. We also determined the phylogenetic composition of the uncultivated Fecal 7 mix sample. The V4 region of the 16S rRNA gene was PCR amplified with dual-indexed Illumina primer-pairs, using different combinations of barcoded forward and reverse primers (v4.SA501-505 and v4.SA701-706, respectively, Table S11) as previously described^46^. The primers consist of the appropriate Illumina adapters, an 8-nt index sequence, a 10-nt pad sequence, a 2-nt linker and specific sites for the V4 region. The PCR reactions consisted of 1.5 μl (30 ng) of template DNA, 10 μl of Phusion HF buffer (Thermo Fisher Scientific), 4 μl of 2.5 mM deoxynucleotide triphosphates mix (dNTPs), 0.5 μl of Phusion DNA polymerase (2 U/μl) (Thermo Fisher Scientific), 1-1 μl of primers, 10 μM each, 3 μl DMSO (100%) and 29 μl of nuclease-free H_2_O in a final reaction volume of 50 μl. The following thermocycler conditions were used: 95 °C for 2 minutes, 25 cycles of 95 °C for 20 seconds + 56 °C for 15 seconds + 72 °C for 5 minutes, and 72 °C for 10 minutes. Following gel electrophoreses of the PCR products, the 400 bp long amplicons were extracted from the gel (Thermo Scientific GeneJET Gel Extraction Kit) and, following a second purification step (Zymo Research DNA Clean & Concentrator™-5 Kit), were sequenced using MiSeq Illumina platform. To prepare the samples for sequencing, the amplicons were quantified using a fluorometric method (Qubit dsDNA BR Assay Kit, Thermo Fischer Scientific) and libraries were mixed with Illumina PhiX in a ratio of 0.95 to 0.05. Sequencing on the Illumina MiSeq instrument was carried out with a v2 500 cycle sequencing kit (Illumina). 100 μM stock custom sequencing primers^46^ were mixed with standard read1, index read and read2 sequencing primers included in the MiSeq cartridge.

After sequencing, 16S rRNA reads were demultiplexed and processed with the *Mothur* software (version 1.36.1, ^47^). The average counts per sample were 21,979. To filter out the low-read-counts we followed the protocol of Rettedal et al. ^26^. The number of sequences per sample was equalized to 20,000 read counts using random re-sampling with a custom *R* script. Reassuringly, 20,000 read counts are well above the threshold where phylogenetic diversities show saturation in the samples (see the rarefaction curves of samples in Figure S11, that were calculated using the *vegan* (version 2.4-3, ^48^) *R* package). Sequences were merged at the level of 97% sequence identity and taxonomically assigned using the Silva ribosomal RNA database^38^. OTUs were classified at the family level since the V4 region allows accurate identification only down to this level^49^ (Table S5). After removal of reads that could not be classified, 362 OTUs remained. To evaluate the reproducibility of the cultivation and sequencing, we generated seven replicates from the untreated samples. Samples referred to as “Untreated 1-5” originate from independent cultivation experiments started from different aliquots of the same five frozen samples (Figure S5, Figure S12). In the case of “Untreated 5, 5i and 5ii” samples cultivations were started from the same sample and cultures were grown in parallel (Figure S5, Figure S12).

To quantify within-sample diversity from 16S rRNA data, we used the *vegan* (version 2.4-3, *R* package^48^ to calculate the most commonly used alpha diversity indices (Shannon index: see Figure 3A; Fisher’s and Inverse Simpson indices: see Figure S6). Unweighted Unifrac distances (Figure 3B) were computed by the *Phyloseq* (version 1.22.3 *R* package^50^).

To identify differentially abundant bacterial families in the resistance microbiota, we applied the *edgeR* (version 3.16.5 *R* package^51^) on the family-level 16S rRNA abundance data of the cultured microbiota samples as suggested earlier ^52^. To this end, abundances were normalized using the TMM method^53^ and then untreated and AMP/AB treated samples were compared using negative binomial tests in a pairwise manner. We used the Benjamini-Hochberg FDR correction method to correct the p-values for multiple testing^54^.

### Comparing the prevalence of AMP- and antibiotic-resistance genes in gut microbial genomes

In order to assess if the large taxonomic diversity in the AMP resistant microbiota corresponds to a diverse reservoir of AMP resistance genes, we compared the prevalence of previously described AMP- and antibiotic-resistance gene families in a representative set of gut microbial genomes as follows. Genome sequences used for the analysis of the mobile gene pool (Table S2) were filtered to retain only those sequences that belong to one of the AMP- and antibiotic-resistant bacterial families (Table S5). In this analysis, we used only the genome sequences from the Human Microbiome Project (HMP) since in the Fiji cohort the family-level taxonomic assignments of the genome sequences were sporadic. In the remaining dataset 24 AMP- and 26 antibiotic-resistant bacterial families were represented with 222 and 219 genome sequences, respectively. To avoid sampling bias due to unequal representation of species among the genome sequences, one genome sequence was randomly picked from each genome group (genomes with lower than 2% 16S rRNA gene dissimilarities, for details, see section *Analysis of the mobile gene pool of the gut microbiota*). Then, for each known AMP and antibiotic resistance gene family, the number of genomes with at least one positive annotation was counted and plotted (Figure S7).

### Comparing phylum-level distributions of the resistant microbiota and the transferring resistance contigs

To compare the phylum-level distributions of resistant gut bacteria and the resistance contigs originating from them (Figure 3D), we carried out logistic regression analyses on count data (Table S9) as follows. For each phylum, we fitted a logistic regression model to predict if a resistant gut microbiota colony or resistance-conferring contig belongs to that particular phylum or to the rest of phyla. Thus, each entry in the dataset represented either a colony from the drug-treated cultivation experiment or a resistance contig detected in the functional metagenomics screen. The predictor variables of the models were i) whether the entry was a resistance contig or a cultivated colony, ii) the type of treatment (AMP or antibiotic) and iii) the interaction of these two variables. As we were interested in whether a given phylum contributed disproportionally more (less) AMP than AB resistance contigs compared to its relative frequency in the drug-treated cultured microbiota, we tested if the interaction term of the logistic regression model was significant (using the *glm* function of *R*).

### Comparison of resistance levels in *E. coli* and Salmonella enterica

In order to investigate whether the level of resistance provided by AMP resistance genes depends more on the genetic background of the recipient bacteria than in the case of antibiotic resistance genes, we measured how DNA fragments that provide AMP or antibiotic resistance to *E. coli* influence susceptibility in a related Enterobacter species, S. *enterica* Serovar Typhimurium LT2. For this purpose, we used a representative set of plasmids carrying resistance DNA fragments that were isolated in our screens from AMP and antibiotic treatments (Table S10). Special care was taken to avoid the inclusion of multiple plasmids carrying resistance genes with the same function or bias toward certain AMPs/ABs. To this end, when it was possible, we selected plasmids from each AMP/AB selection in equal numbers carrying resistance genes with different functions. This resulted in 16 plasmids and 25 AMPs and ABs. The provided resistance levels (MIC fold changes) were measured for both species with standard micro-dilution assay the same way as described in the section below.

### Quantification of the resistance gains that metagenomic contigs provide

To investigate the resistance gains that contigs provide for the recipient bacteria against AMP and antibiotic treatments, we carried out minimum inhibitory concentration (MIC) measurements with selected contigs from the uncultured and cultured microbiota. The resistance gains were quantified by measuring minimum inhibitory concentrations (MIC) with a standard broth microdilution method^41^. Briefly, 5000 *E. coli S. enterica* cells grown overnight in MS medium with 50 μg/ml kanamycin were used to inoculate wells of a 96-well plate. 3 rows on the plate were inoculated with the strain in question and 3 rows with control cells. As a control, *E. coli* BW25113 or S. *enterica* Serovar Typhimurium LT2 strains carrying the control plasmid was used as in the functional metagenomic selection experiments. Prior to inoculation, each well on the plate was prefilled with 100 μl modified MS medium supplemented with the proper concentration of AMP or AB. On the plate each AMP/AB was represented in 11 different concentrations (3 wells /concentration / clone or control). 3 wells contained only medium to check the growth in the absence of any antimicrobial. MIC values were determined by measuring OD_600_ after 16-24 hours of incubation at 30°C with a continuous shaking at 240 rpm.

### Functional analysis of a putative LpxF from the metagenomic selections

The aim of the functional characterization was twofold. First, to support the functional prediction for the identified LpxF orthologs with a biochemical assay. Second, to estimate quantitatively the extent to which these LpxF orthologs reduce the net negative surface charge of the bacterial cell as compared to a previously characterized LpxF from *Francisella novicida* that removes >90 % of the 4’-phosphate groups from the pentaacylated lipid A molecules and hence alters the charge of the outer membrane^55^. To estimate the surface charge of bacterial cells, we used a fluorescein isothiocyanate (FITC)-labeled poly-L-lysine (PLL) (FITC-PLL) (Sigma-Aldrich^®^) based assay. Poly-L-lysine is a polycationic molecule, widely applied to study the interaction between charged bilayer membranes and cationic peptides^56^. The following strains were used in this measurement: *E. coli* BW25113 Δ*lpxM, E. coli* BW25113 Δ*lpxM* carrying the pZErO-2 plasmid with the LpxF ortholog from *Parabacteroides merdae* ATCC 43184 identified in our selection experiments (Table 1), *E. coli* BW25113 Δ*lpxM* carrying the pWSK29 plasmid with LpxF from *Francisella novicida* (*53*) and *Bacteroides thetaiotaomicron*. We measured the phenotypic effect of both LpxF carrying plasmids on Δ*lpxM* genetic background, since LpxF from *F. novicida* cannot carry out its biological function in wild-type *E. coli*, only when the lipid A molecules are tetra- or pentaacylated as it is in the case of Δ*lpxM E. coli*^55^. *B. thetaiotaomicron* intrinsically expresses an *lpxF* ortholog (BT1854) which is responsible for the high level of resistance of this strain against Polymyxin B^12^. Prior to the surface charge measurements, cells were grown overnight in TYG (Tryptone Yeast Extract Glucose) medium^57^ in anaerobic conditions. We grew all the strains in TYG medium to allow the comparability with *B. thetaiotaomicron*. Cells were washed twice with phosphate-buffered saline (PBS) then resuspended to a cell density of OD_600_ = 1. Cells were incubated with 2 μl of 5 mg/ml FITC-PLL and 100 μl of 1 μg/ml 4,6-diamidino-2-phenylindole (DAPI) for 10 minutes, followed by centrifugation (4500 rpm, 5’). DAPI was used in order to identify the live cells. Cells were washed twice with PBS then diluted 100-fold in 100 μl of PBS and transferred into a black clear-bottom 96-well microplate (Greiner Bio-One). Prior to fluorescent microscopy analysis, cells were collected to the bottom of the plate by centrifugation (4500 rpm, 10’). Pictures were taken with a PerkinElmer Operetta microscope using a 60x high-NA objective to visualize the cells. Images of two channels (DAPI and FITC-PLL) were collected from ten sites of each well. Mean fluorescent intensity for each well was calculated using the Harmony^®^ High Content Imaging and Analysis Software. Experiments were carried out in 4 biological replicates.

---

**Figure S1.**
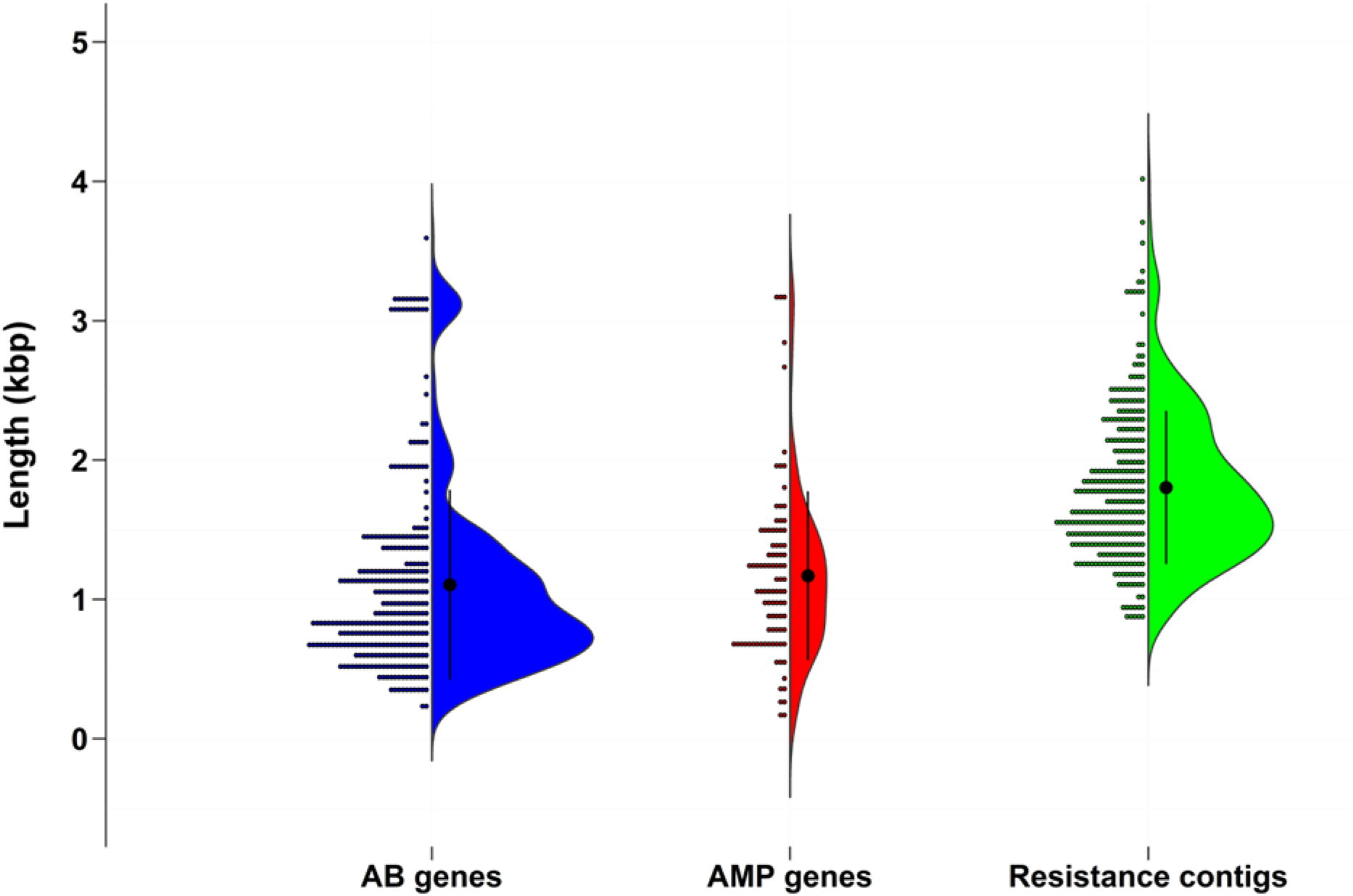
The length distributions of known antibiotic- and AMP-resistance genes are well within the fragment size range of the metagenomic library. Blue and red plots show the length (kilobase pair (kbp)) distribution of known antibiotic- and AMP-resistance genes, respectively, from a representative set of gut microbial genomes (Table S2). Green plot shows the length (kbp) distribution of all antibiotic- and AMP-resistance DNA fragments identified in our functional metagenomic selections (Table S4). The mean value and standard deviation are shown on the plots.

**Figure S2.**
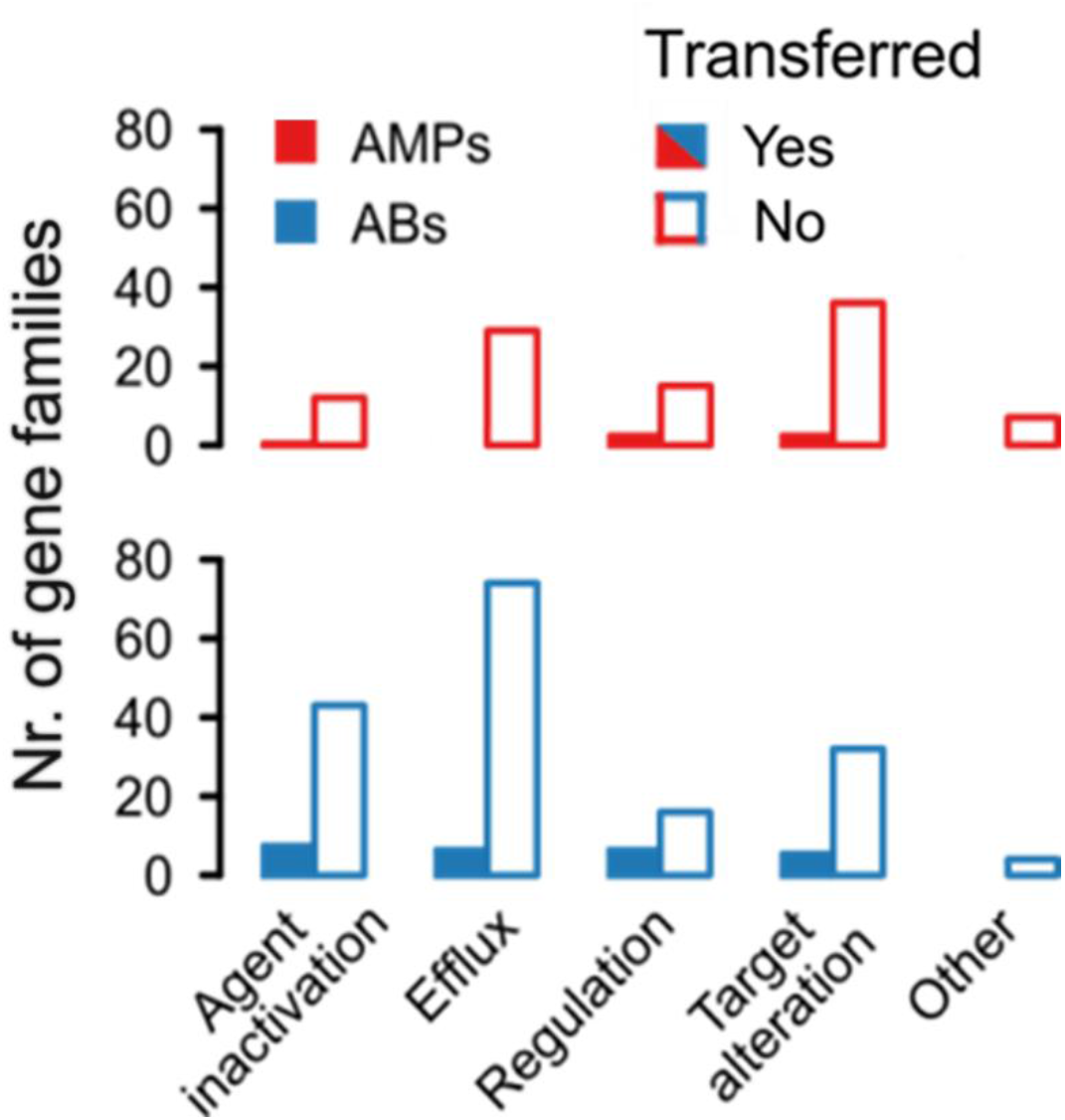
Presence (filled bars) / absence (empty bars) patterns of the known AMP- and antibiotic-resistance gene families on the metagenomic contigs identified in our functional selections. Genes were assigned to gene families (orthogroups) which were classified into major functional categories (see Methods). A gene family was considered present if at least one resistance gene from its orthogroup had a significant sequence similarity hit on the DNA fragments (see Methods). We considered a resistance gene family absent if the orthogroup did not result in any hits on the metagenomic contigs from the functional selections. *n*=200 for ABs, *n*=105 for AMPs.

**Figure S3.**
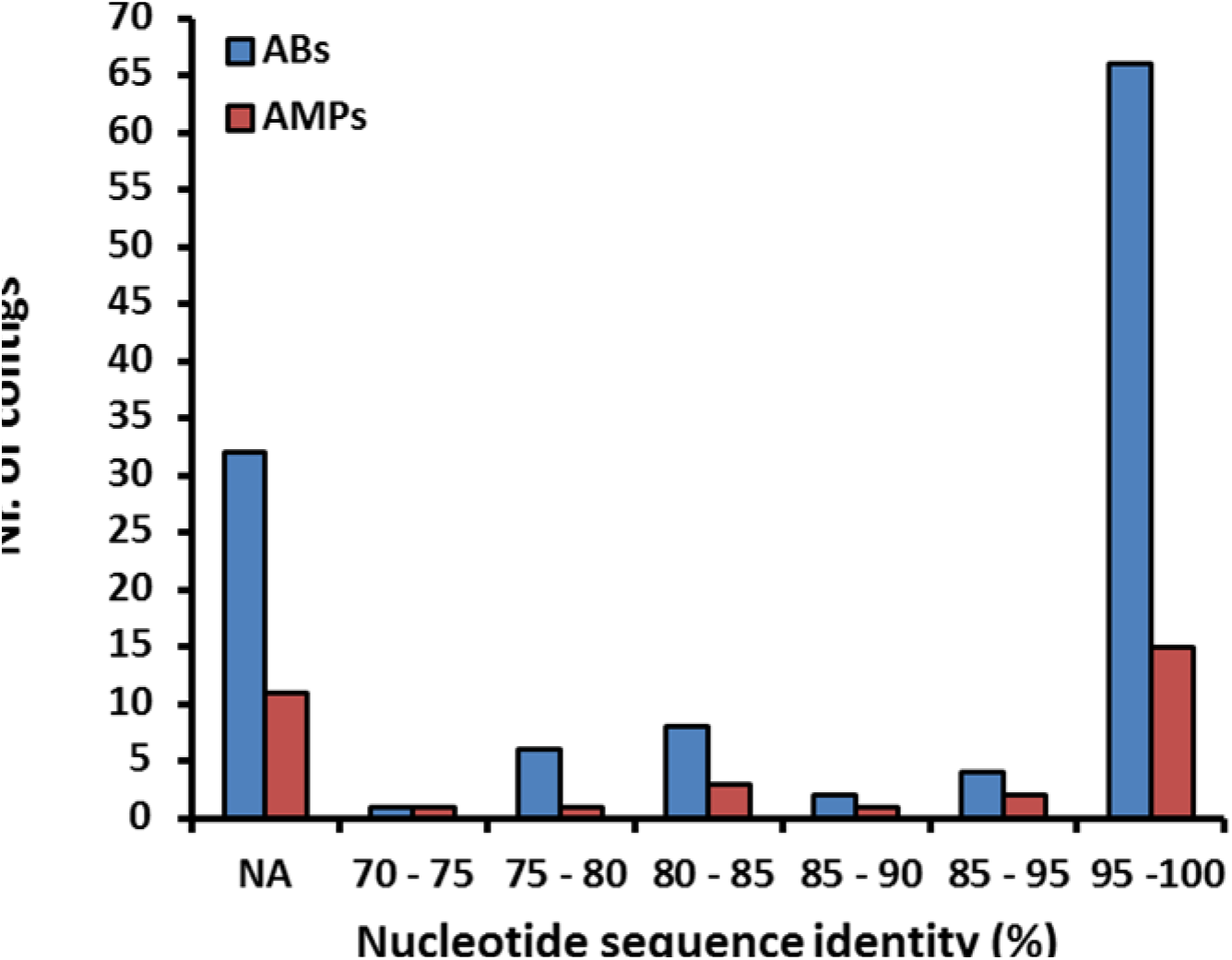
Distribution of the nucleotide sequence identities between the AMP/AB resistance contigs originating from the metagenomic libraries of the uncultured microbiota (Table S4) and the genome sequences from the Human Microbiome Project (see Methods). 28 % of the contigs (marked with NA) did not result in significant alignment to the genome sequences in the HMP database. *n*=119 for ABs, *n*=33 for AMPs.

**Figure S4.**
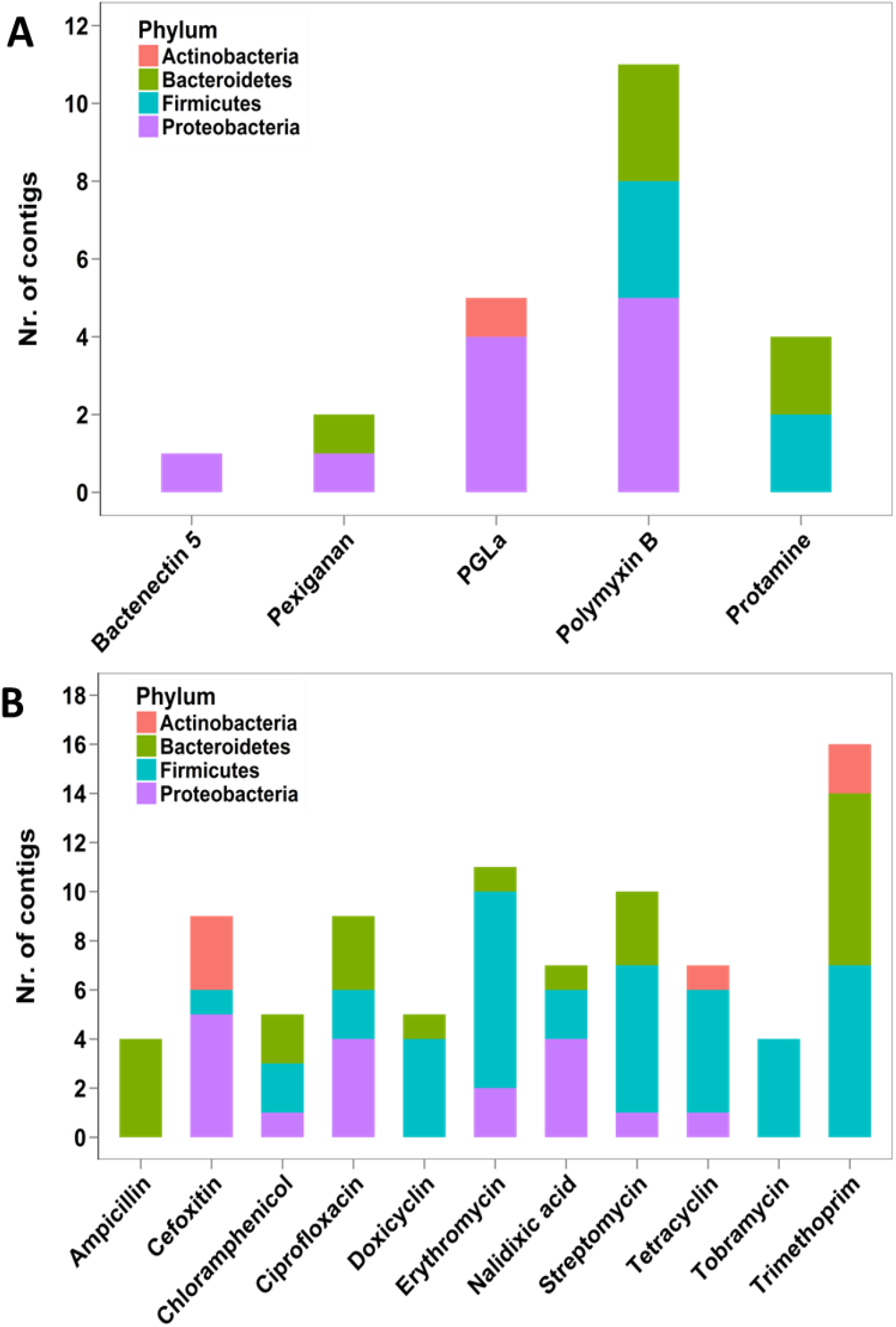
Phylum-level distribution of resistance contigs originating from the functional selection of the uncultured microbiota with different A) AMPs (*n*=23) and B) antibiotics (*n*=87) (Table S4).

**Figure S5.**
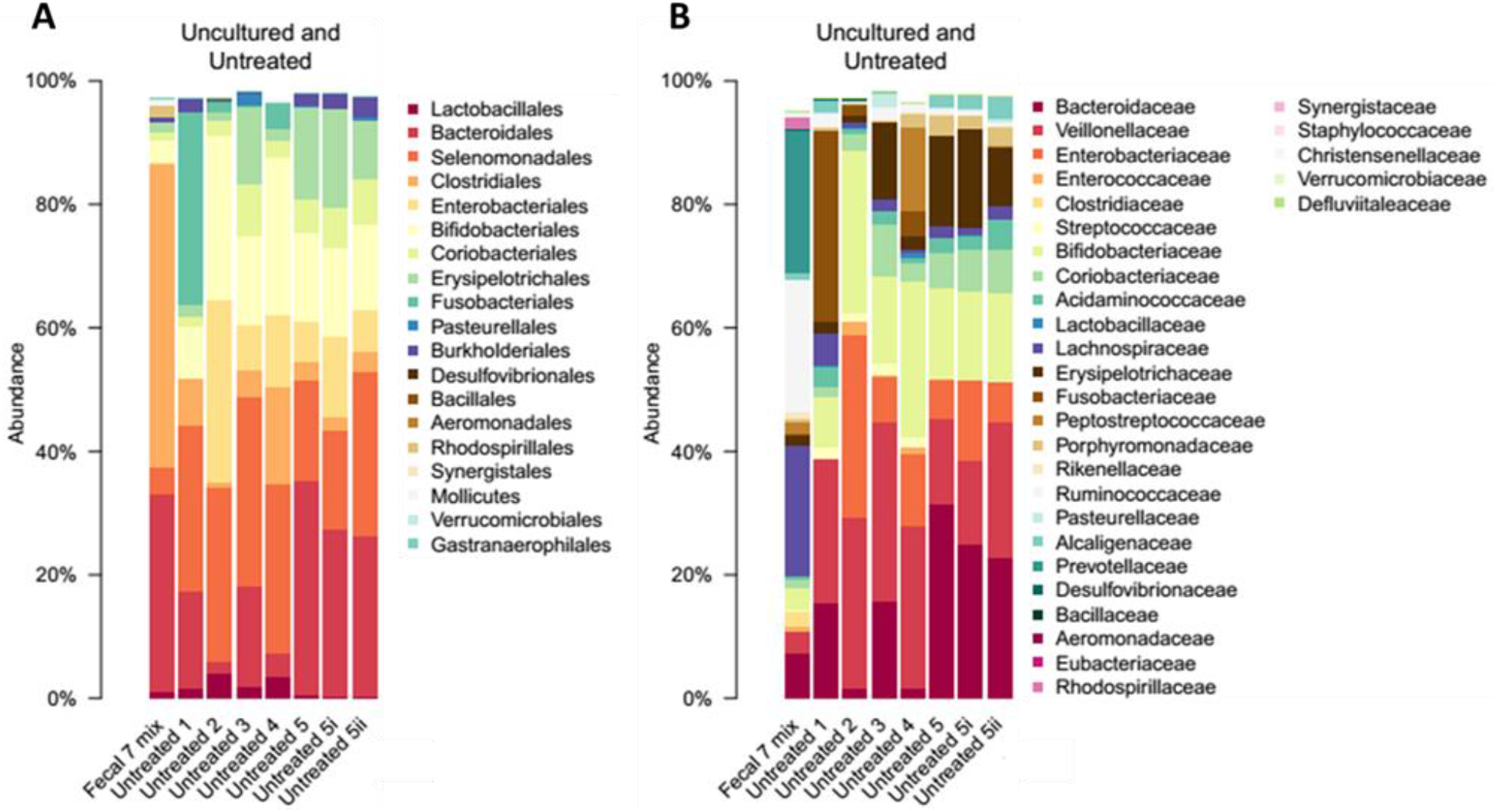
Percent abundances of bacterial orders (A) and families (B) in the uncultured (Fecal 7 mix) and anaerobically cultured (Untreated 1-5, 5i, 5ii) gut microbial samples from seven unrelated healthy individuals. At order level, the cultivable proportion was 64-86 %, while at the family level it was 65-74 %. These results are consistent with the cultivation efficiency reported in a previous study^26^. “Untreated 1-5” samples are biological replicates started from different aliquots of the same frozen samples in independent cultivation experiments. “Untreated 5i” and “Untreated 5ii” are technical replicates of the “Untreated 5” sample started from the same sample and grown at the same time in the same experiment. Data is presented in Table S5.

**Figure S6.**
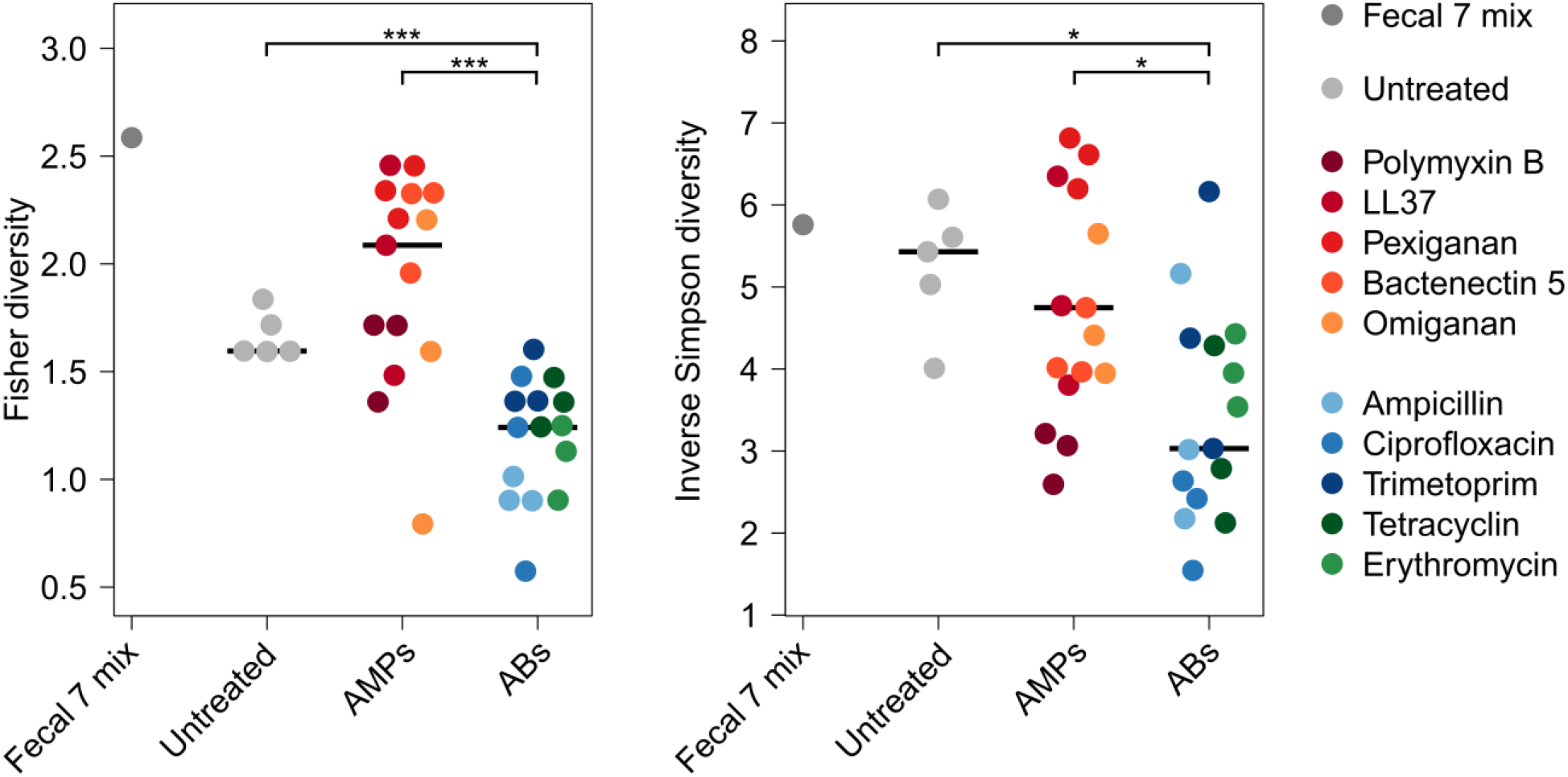
Diversities of the cultured microbiota and the original faecal sample (Fecal 7 mix). Fisher (A)^58^ and Inverse Simpson (B)^59^ alpha diversity indices were calculated at the bacterial family level. Alpha diversity reflects the number of different taxa and distribution of abundances. Asterisks indicate significant difference, Mann-Whitney U tests: * indicates *P* < 0.05, *** indicates *P* < 0.001, *n*=36. Central horizontal bars represent median values.

**Figure S7.**
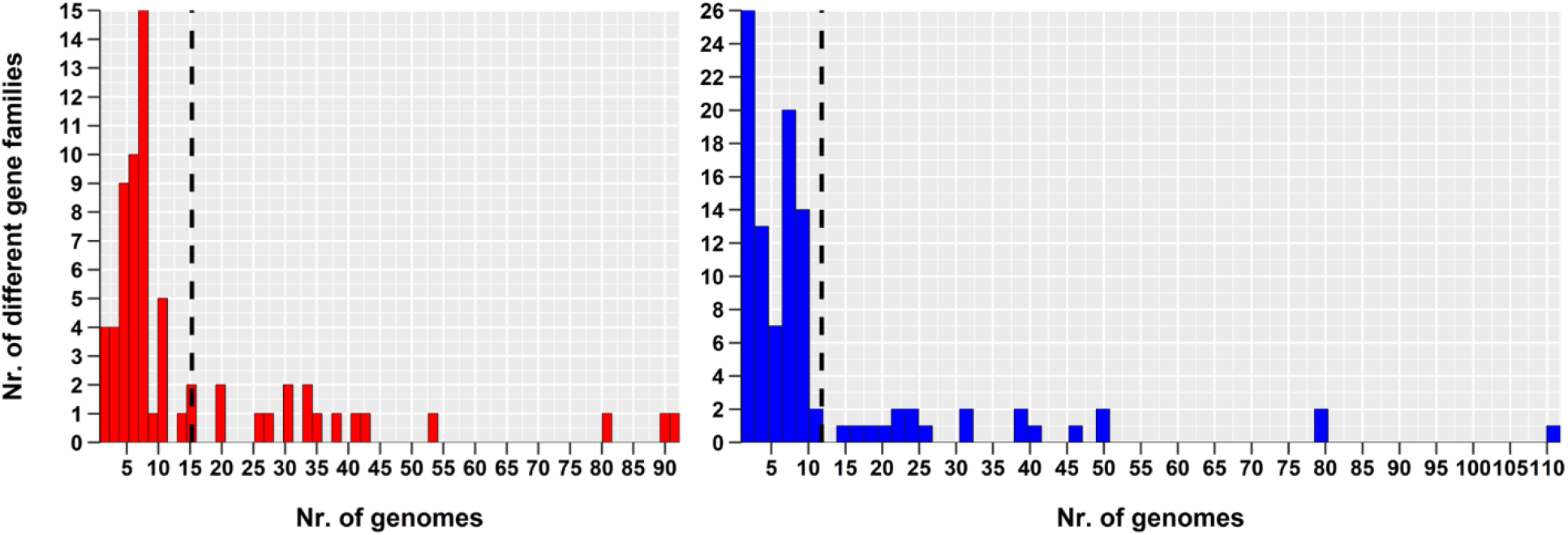
The diversity of previously described AMP-(red) and antibiotic resistance genes (blue) in a representative set of gut microbial genomes^15^. The dashed line represents the mean value. For details see the section “*Comparing the prevalence of AMP- and antibiotic-resistance genes in gut microbial genomes*”. (P<10^-16^, two-sided negative binomial regression, *n*=67 (AMPs) and *n*=102 (ABs)).

**Figure S8.**
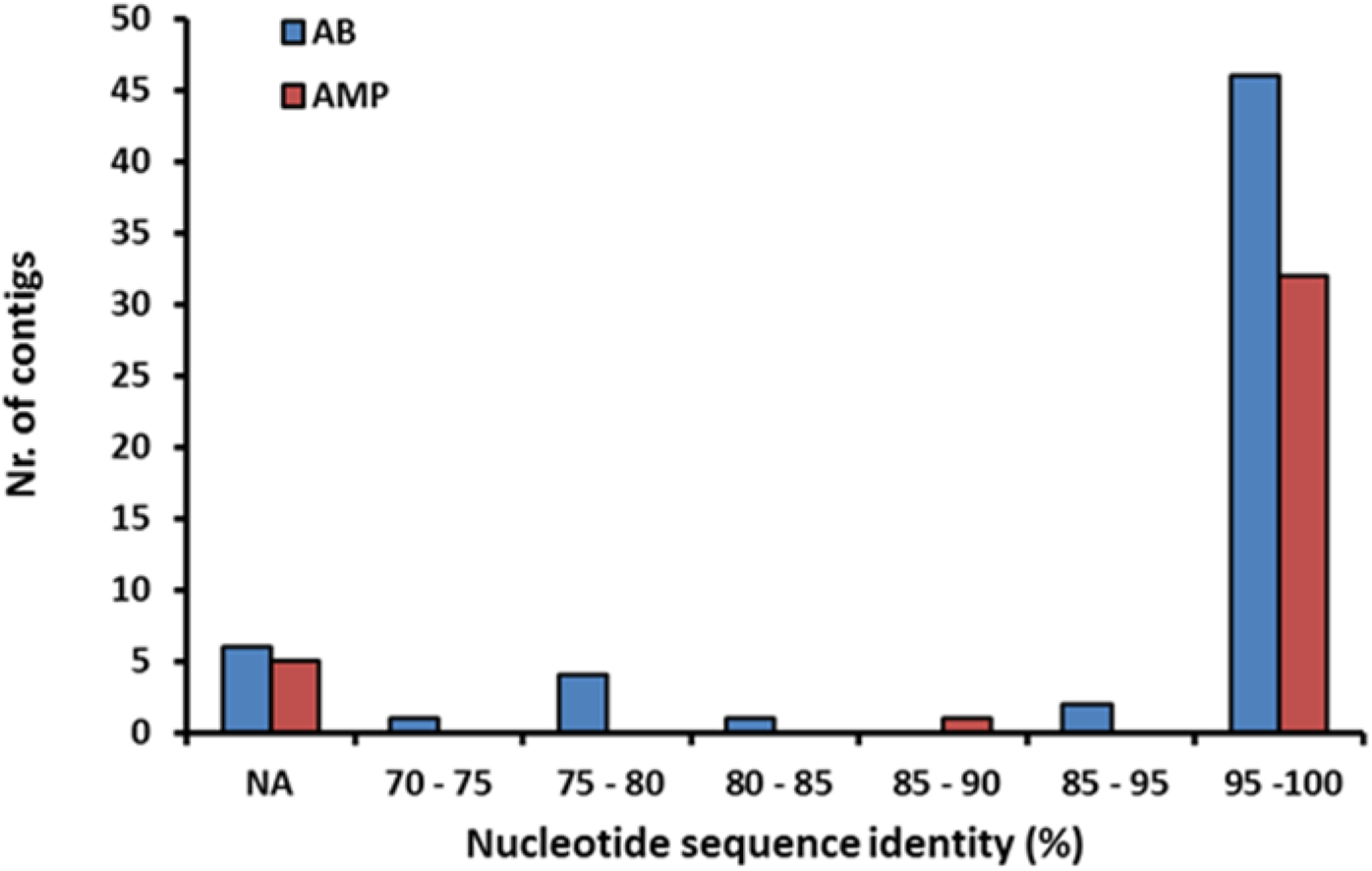
Distributions of the nucleotide sequence identities between the AMP/AB resistance contigs originating from the cultured microbiota and the genome sequences from the Human Microbiome Project (see Methods), *n*=60 for ABs, *n*=38 for AMPs (Table S8). 11 % of the contigs (marked with NA) did not result in significant alignment to the genome sequences in the HMP database.

**Figure S9.**
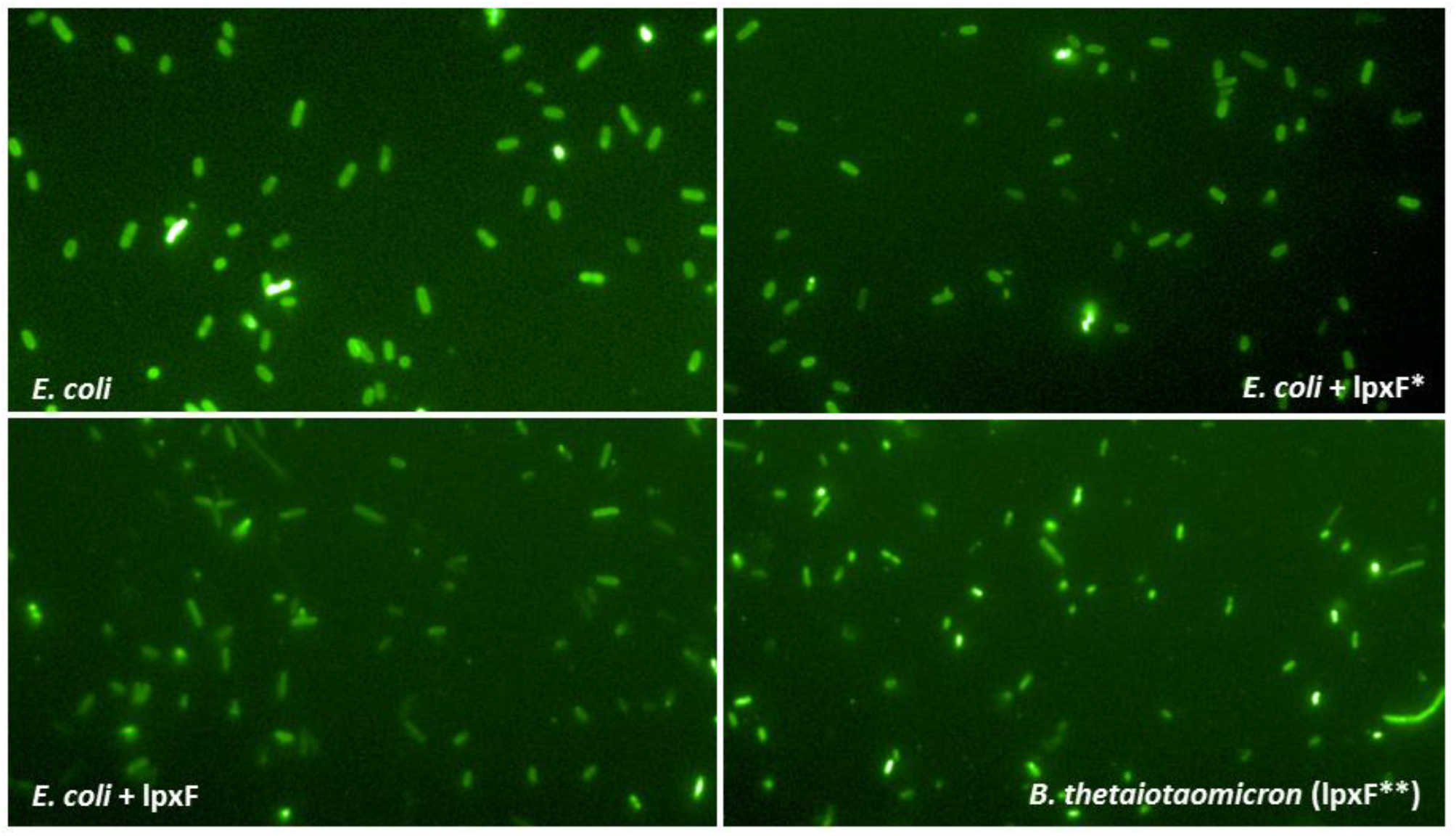
Microscopic images of cells incubated with FITC-labeled poly-L-lysine (PLL) polycationic molecules. The brighter are the cells the more PLL is bound, indicating a more negative outer membrane surface charge. For details see “Surface charge measurement” section in Supplementary methods. The quantitative analysis of the fluorescent signals is presented in Figure 4B. Abbreviations: *E. coli* = BW25113 Δ*lpxM*^¥^. *E. coli* + *lpxF* = *E. coli* expressing an ortholog gene of *lpxF* from *Parabacteroides merdae* ATCC 43184. This gene was identified in our metagenomic screen (see Table 1). *E. coli* + *lpxF*^*^ = *E. coli* expressing the *LpxF* of *Francisella tularensis subsp. novicida. B. thetaiotaomicron* (*lpxF*^**^) = *Bacteroides thetaiotaomicron* strain, which intrinsically expresses the *lpxF BT1854* gene. ^¥^ In BW25113 Δ*lpxM* strain the *lpxM* gene is deleted and therefore this strain has pentaacylated lipid-A molecules instead of the hexaacylated ones^60^. We used this strain to allow the phenotypic comparison of LpxF to the *LpxF* from *Francisella tularensis*, which can be expressed only in ΔlpxM *E. coli* background that has pentaacylated LPS molecules (*n*=4).

**Figure S10.**
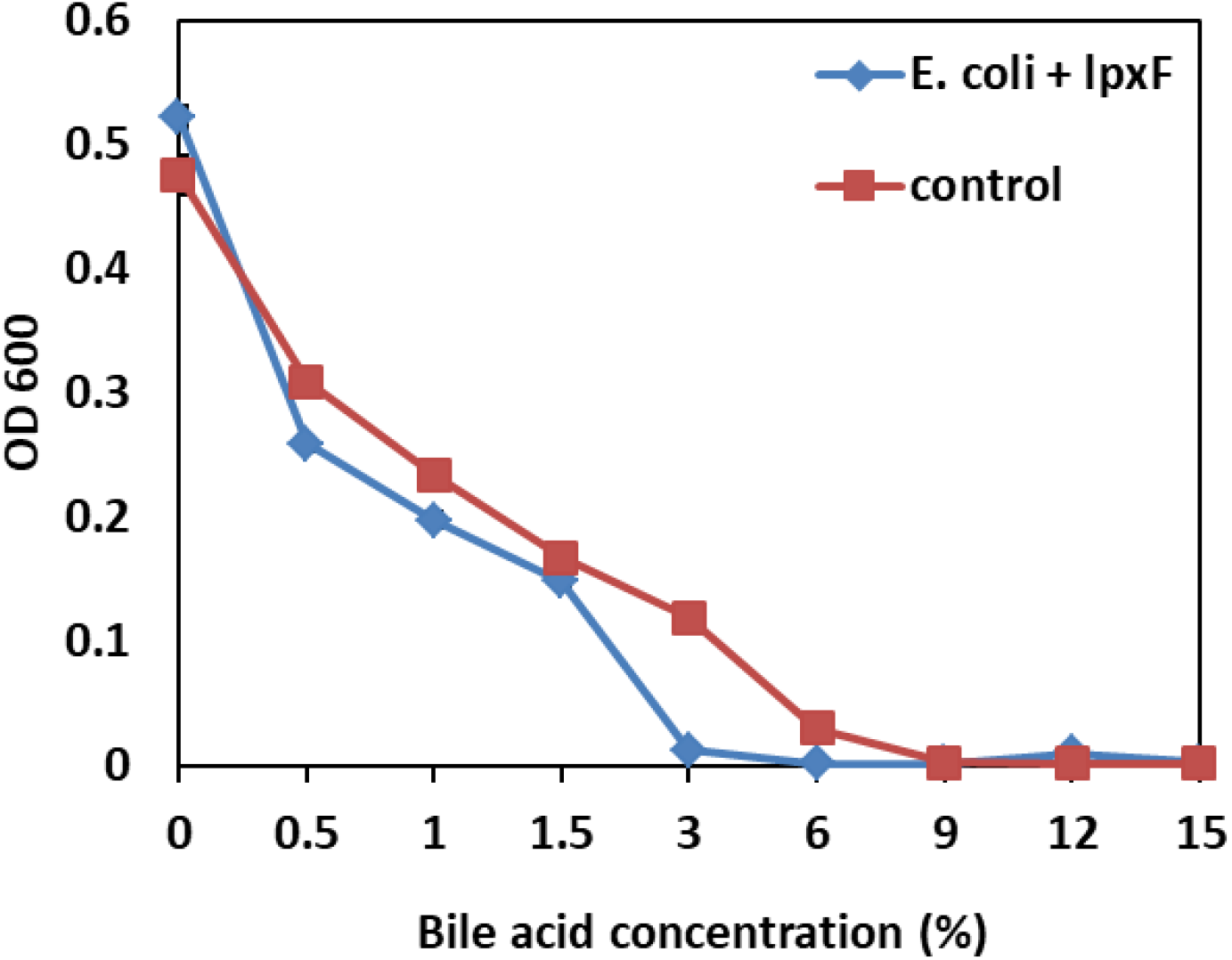
The expression of the *lpxF* ortholog gene from *Parabacteroides merdae ATCC 43184* (see Table 1.) increases the sensitivity of *E.coli* to bile acid, a membrane-damaging agent. Abbreviations: E. coli + lpxF - *E. coli* BW25113 expressing the *lpxF* ortholog gene from *Parabacteroides merdae ATCC 43184* identified during the functional selection of the metagenomic libraries against polymyxin B, control - *E. coli* BW25113 harbouring the library vector with a random metagenomic DNA insert which does not influence resistance against AMPs or antibiotics (*n*=3).

**Figure S11.**
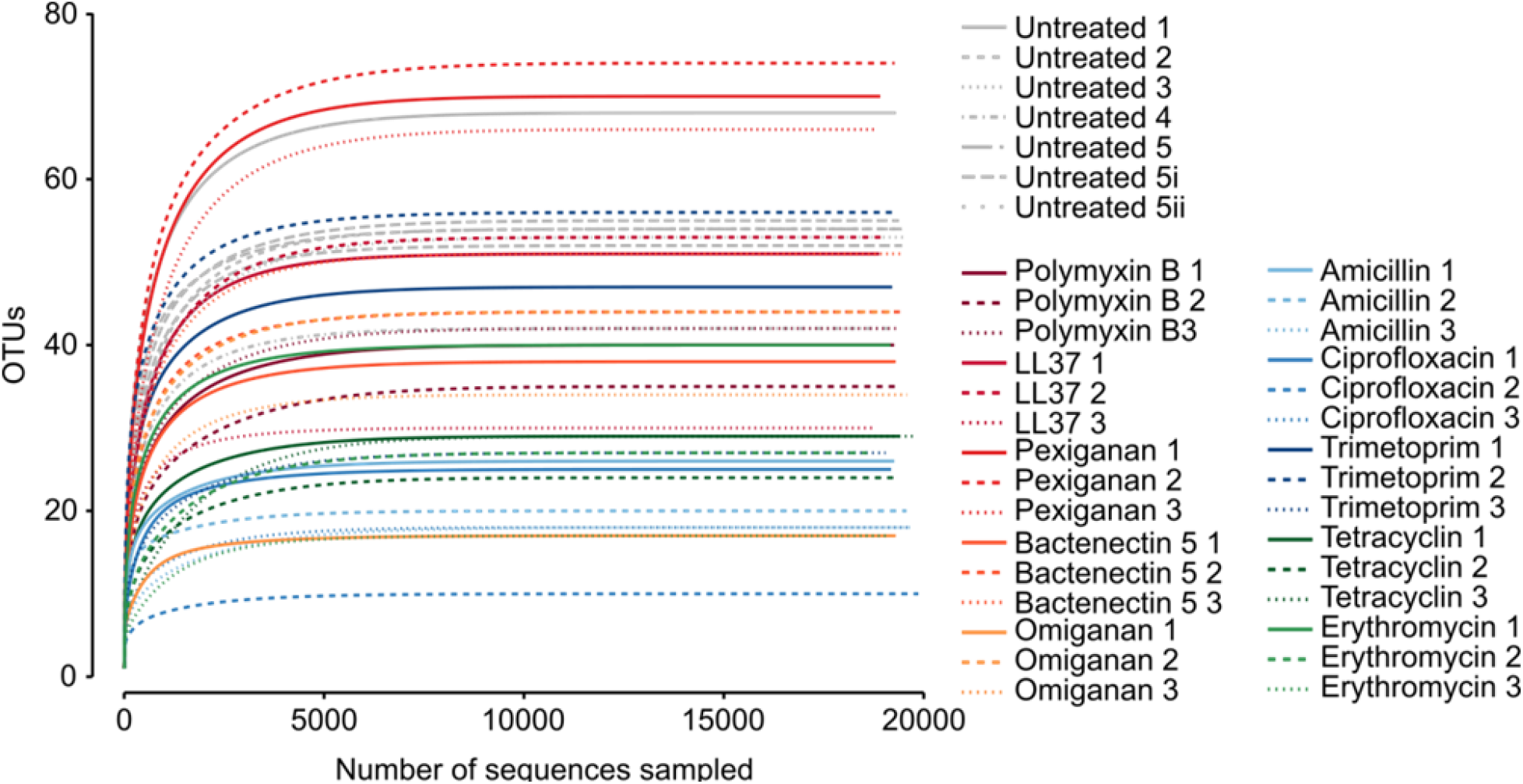
Rarefaction OTU (‘species’) richness curves. Curves are plots of the number of OTUs as a function of the number of sequences, for different samples. At a sample size of 10,000 sequences, all curves reach their plateau, indicating that sufficient sequences were used to estimate the number of OTUs in different samples.

**Figure S12.**
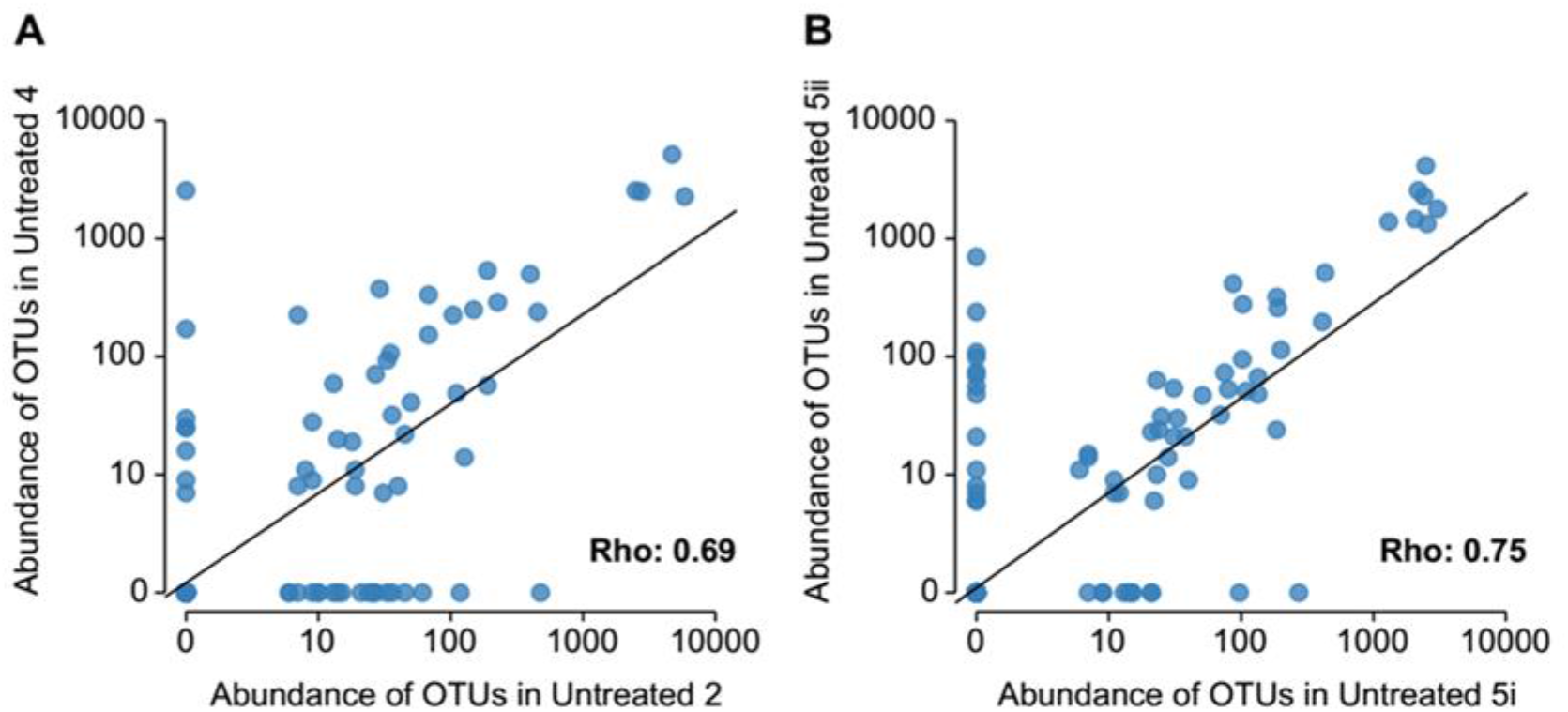
OTUs abundances correlate across biological (A) and technical (B) replicates of cultivated, untreated samples, showing good reproducibility of the replicates. Spearmann’s rho is 0.69 (A) and 0.79 (B), respectively. Points represent individual OTUs. *n*=2, Untreated 2 and 4 samples are biological replicates, while Untreated 5i and 5ii are technical replicates. Biological replicates originate from independent cultivation experiments starting from different aliquots of the same frozen samples. Technical replicates originate from parallel cultivations starting from the same sample and grown at the same time.

**Table S1**. A comprehensive catalogue of previously reported AMP resistance genes compiled based on literature mining and of antibiotic resistance genes from the CARD database^35^. The table is provided as a separate Excel file.

**Table S2**. Identification of the AMP- and antibiotic-resistance genes in the mobile gene pool and in the genomes from which the mobile gene pool was derived. The table is provided as a separate Excel file.

**Table S3.** List and characteristics of antimicrobials used in this study. The origin of the antimicrobial peptides is indicated in brackets and the peptides marked with asterisks are involved in clinical trials.

| Antimicrobials | Mode of action | Used in |
| --- | --- | --- |
| **<u>ANTIMICROBIAL PEPTIDES (AMPS)</u>** |  |  |
| **Cecropin P1 (parasitic nematode)** | pore-forming activity | Functional metagenomics, culturing the gut microbiota |
| **Bactenectin 5 (cow)** | intracellular targets | Functional metagenomics, culturing the gut microbiota |
| **Indolicidin (cow)** | pore-forming activity, inhibition of DNA synthesis | Functional metagenomics |
| **LL37 (human)** | pore-forming activity, membrane depolarization | Functional metagenomics, culturing the gut microbiota |
| **Omiganan\* (synthetic, indolicidin analogue)** | pore-forming activity, inhibition of DNA synthesis | Functional metagenomics, culturing the gut microbiota |
| **Pexiganan\* (synthetic, magainin analogue)** | pore-forming activity | Functional metagenomics, culturing the gut microbiota |
| **PGLa (frog)** | pore-forming activity | Functional metagenomics |
| **Polymyxin B (bacterial)** | pore-forming activity, membrane depolarization | Functional metagenomics, culturing the gut microbiota |
| **PR-39 (pig)** | inhibition of DNA and protein synthesis | Functional metagenomics |
| **Protamine (salmon)** | intracellular targets | Functional metagenomics |
| **R8 (synthetic)** | pore-forming activity | Functional metagenomics |
| **Tachiplesin II (horseshoe crab)** | pore-forming activity, membrane depolarization | Functional metagenomics |
| **<u>ANTIBIOTICS (ABS)</u>** |  |  |
| **Ampicillin** | inhibition of cell wall synthesis | Functional metagenomics, culturing the gut microbiota |
| **Cefoxitin** | inhibition of cell wall synthesis | Functional metagenomics |
| **Ciprofloxacin** | inhibition of DNA synthesis | Functional metagenomics, culturing the gut microbiota |
| **Chloramphenicol** | inhibition of protein synthesis | Functional metagenomics |
| **Doxycycline** | inhibition of protein synthesis | Functional metagenomics |
| **Erythromycin** | inhibition of protein synthesis | Functional metagenomics, culturing the gut microbiota |
| **Nalidixic acid** | inhibition of DNA synthesis | Functional metagenomics |
| **Streptomycin** | inhibition of protein synthesis | Functional metagenomics |
| **Tetracycline** | inhibition of protein synthesis | Functional metagenomics, culturing the gut microbiota |
| **Tobramycin** | inhibition of protein synthesis | Functional metagenomics |
| **Trimethoprim** | inhibition of folate synthesis | Functional metagenomics, culturing the gut microbiota |

**Table S4**. List of resistance contigs identified from the functional metagenomics selections of the uncultured microbiota with 12 AMPs and 11 small-molecule antibiotics (Table S3). The table is provided as a separate Excel file.

**Table S5**. Bacterial abundances at family and order levels in the untreated and AMP/antibiotic resistant microbiota. Abundances at order and family levels are presented on separate Excel sheets. The table is provided as a separate Excel file.

**Table S6.** Selection conditions used for the anaerobic cultivation experiments. Those AMP/AB treatment concentrations were chosen where the average proportion of the resistant colonies originating from three selection concentrations per AMP/AB ranged from 0.01 to 0.1 % of the total cultivable colony number in the absence of any antimicrobial. Trimethoprim represented an exception, where the resistant fraction was slightly higher, but the concentration could not be raised further because of solubility issues.

| Antimicrobial peptide (AMP) / antibiotic (AB) | Selection concentration (µg/ml) | Treatment type | Relative resistance level of the gut microbiota* | Resistant colonies (percentage of the cultivable cell population) |
| --- | --- | --- | --- | --- |
| Bactenectin 5 | 666.7 | AMP | 6.7 | 0.02 |
| LL37 | 1400 | AMP | 10 | 0.02 |
| Omiganan | 250 | AMP | 8.3 | 0.04 |
| Pexiganan | 166.7 | AMP | 16.7 | 0.01 |
| Polymyxin B | 280 | AMP | 1400 | 0.1 |
| Ampicillin | 700 | AB | 46.7 | 0.1 |
| Ciprofloxacin | 75 | AB | 833.3 | 0.03 |
| Erythromycin | 1600 | AB | 40 | 0.04 |
| Tetracycline | 48 | AB | 26.7 | 0.1 |
| Trimetoprim | 1200 | AB | 1500 | 0.49 |

**Table S7.**
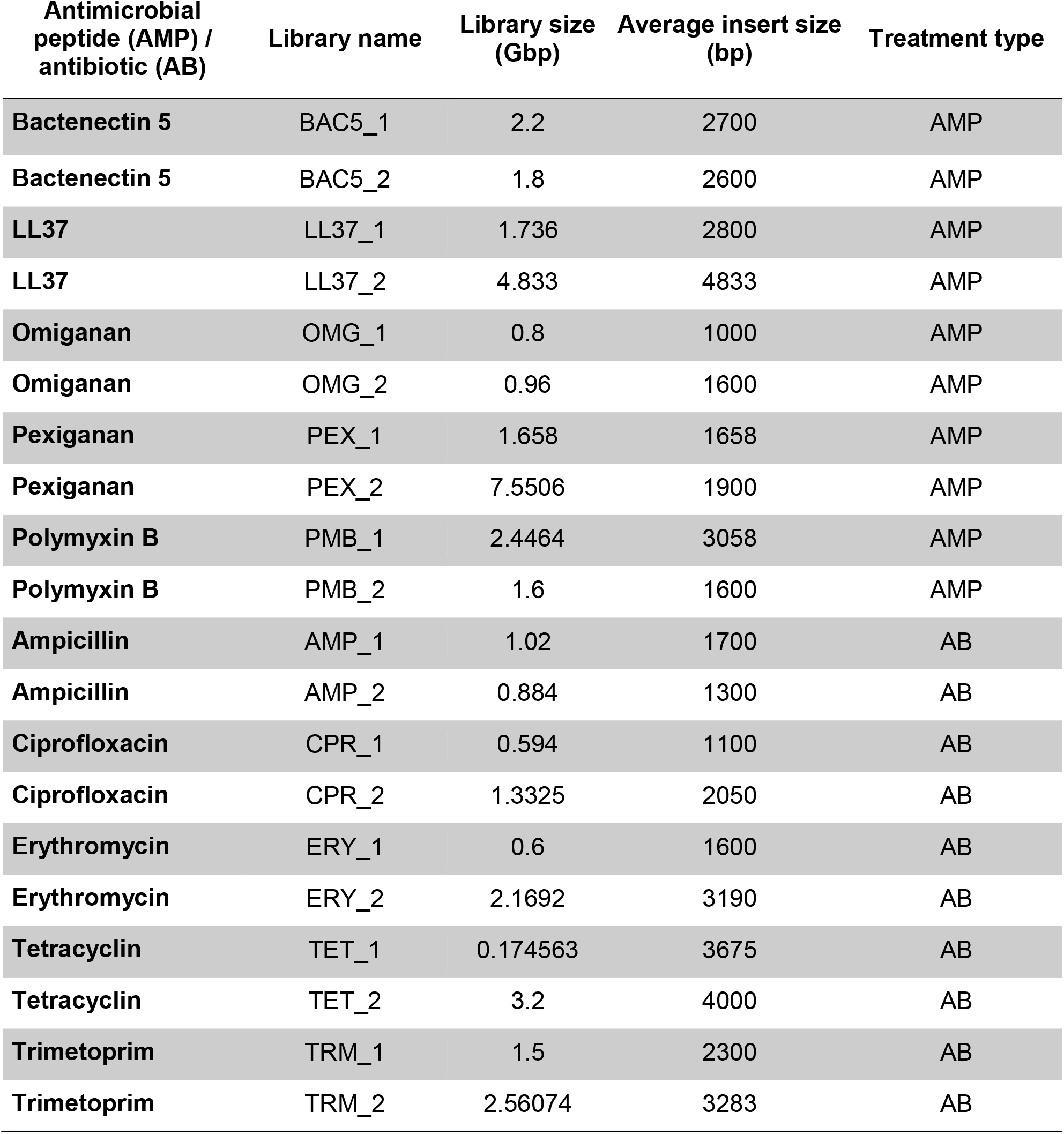
Characteristics of metagenomic libraries constructed from AMP- and antibiotic-resistant microbiota cultures with established microbial compositions.

| Antimicrobial peptide (AMP) / antibiotic (AB) | Library name | Library size (Gbp) | Average insert size (bp) | Treatment type |
| --- | --- | --- | --- | --- |
| Bactenectin 5 | BAC5_1 | 2.2 | 2700 | AMP |
| Bactenectin 5 | BAC5_2 | 1.8 | 2600 | AMP |
| LL37 | LL37_1 | 1.736 | 2800 | AMP |
| LL37 | LL37_2 | 4.833 | 4833 | AMP |
| Omiganan | OMG_1 | 0.8 | 1000 | AMP |
| Omiganan | OMG_2 | 0.96 | 1600 | AMP |
| Pexiganan | PEX_1 | 1.658 | 1658 | AMP |
| Pexiganan | PEX_2 | 7.5506 | 1900 | AMP |
| Polymyxin B | PMB_1 | 2.4464 | 3058 | AMP |
| Polymyxin B | PMB_2 | 1.6 | 1600 | AMP |
| Ampicillin | AMP_1 | 1.02 | 1700 | AB |
| Ampicillin | AMP_2 | 0.884 | 1300 | AB |
| Ciprofloxacin | CPR_1 | 0.594 | 1100 | AB |
| Ciprofloxacin | CPR_2 | 1.3325 | 2050 | AB |
| Erythromycin | ERY_1 | 0.6 | 1600 | AB |
| Erythromycin | ERY_2 | 2.1692 | 3190 | AB |
| Tetracyclin | TET_1 | 0.174563 | 3675 | AB |
| Tetracyclin | TET_2 | 3.2 | 4000 | AB |
| Trimetoprim | TRM_1 | 1.5 | 2300 | AB |
| Trimetoprim | TRM_2 | 2.56074 | 3283 | AB |

**Table S8**. List of resistance contigs identified from the functional selection of the cultured microbiota with 5 AMPs and 5 small-molecule antibiotics (Table S3). The table is provided as a separate Excel file.

**Table S9.** Phylum-level distributions of the AMP- and antibiotic-resistant microbiota and the transferring resistance contigs originating from them. Count data represent either colony numbers from the culturing experiments estimated by using the 16S rRNA abundance data (Table S5) or the number of resistance contigs detected in the functional metagenomics screens. The data was used for logistic regression analyses to determine if the phylum-level representation of resistance contigs are proportional to that of the cultured microbiota (see Methods).

| Phylum | Source | Treatment | Count |
| --- | --- | --- | --- |
| **Proteobacteria** | Transferred DNA contig | AMP | 18 |
| **Proteobacteria** | Transferred DNA contig | AB | 3 |
| **Proteobacteria** | Gut bacteria colony number | AMP | 5590 |
| **Proteobacteria** | Gut bacteria colony number | AB | 1838 |
| **Firmicutes** | Transferred DNA contig | AMP | 5 |
| **Firmicutes** | Transferred DNA contig | AB | 17 |
| **Firmicutes** | Gut bacteria colony number | AMP | 14734 |
| **Firmicutes** | Gut bacteria colony number | AB | 11402 |
| **Bacteroidetes** | Transferred DNA contig | AMP | 8 |
| **Bacteroidetes** | Transferred DNA contig | AB | 30 |
| **Bacteroidetes** | Gut bacteria colony number | AMP | 1050 |
| **Bacteroidetes** | Gut bacteria colony number | AB | 8077 |
| **Actinobacteria** | Transferred DNA contig | AMP | 0 |
| **Actinobacteria** | Transferred DNA contig | AB | 4 |
| **Actinobacteria** | Gut bacteria colony number | AMP | 647 |
| **Actinobacteria** | Gut bacteria colony number | AB | 2868 |
| **Fusobacteria** | Transferred DNA contig | AMP | 0 |
| **Fusobacteria** | Transferred DNA contig | AB | 0 |
| **Fusobacteria** | Gut bacteria colony number | AMP | 630 |
| **Fusobacteria** | Gut bacteria colony number | AB | 151 |

**Table S10**. Resistance gains in *E. coli* and S. *enterica* provided by a representative set of plasmids carrying AMP/antibiotic resistance contigs that were isolated in our metagenomic screens. The table is provided as a separate Excel file.

**Table S11**. List of primers used in this study. The table is provided as a separate Excel file.

